# Ferroptosis regulates hemolysis in stored murine and human red blood cells

**DOI:** 10.1101/2024.06.11.598512

**Authors:** Angelo D’Alessandro, Gregory R. Keele, Ariel Hay, Travis Nemkov, Eric J. Earley, Daniel Stephenson, Matthew Vincent, Xutao Deng, Mars Stone, Monika Dzieciatkowska, Kirk C. Hansen, Steve Kleinman, Steven L Spitalnik, Nareg H. Roubinian, Philip J Norris, Michael P. Busch, Grier P Page, Brent R. Stockwell, Gary A. Churchill, James C Zimring

## Abstract

Red blood cell (RBC) metabolism regulates hemolysis during aging in vivo and in the blood bank. Here, we leveraged a diversity outbred mouse population to map the genetic drivers of fresh/stored RBC metabolism and extravascular hemolysis upon storage and transfusion in 350 mice. We identify the ferrireductase Steap3 as a critical regulator of a ferroptosis-like process of lipid peroxidation. Steap3 polymorphisms were associated with RBC iron content, in vitro hemolysis, and in vivo extravascular hemolysis both in mice and 13,091 blood donors from the Recipient Epidemiology and Donor evaluation Study. Using metabolite Quantitative Trait Loci analyses, we identified a network of gene products (FADS1/2, EPHX2 and LPCAT3) - enriched in donors of African descent - associated with oxylipin metabolism in stored human RBCs and related to Steap3 or its transcriptional regulator, the tumor protein TP53. Genetic variants were associated with lower in vivo hemolysis in thousands of single-unit transfusion recipients.

**Highlights:**

- Steap3 regulates lipid peroxidation and extravascular hemolysis in 350 diversity outbred mice
- Steap3 SNPs are linked to RBC iron, hemolysis, vesiculation in 13,091 blood donors
- mQTL analyses of oxylipins identified ferroptosis-related gene products FADS1/2, EPHX2, LPCAT3
- Ferroptosis markers are linked to hemoglobin increments in transfusion recipients

**Graphical abstract:** 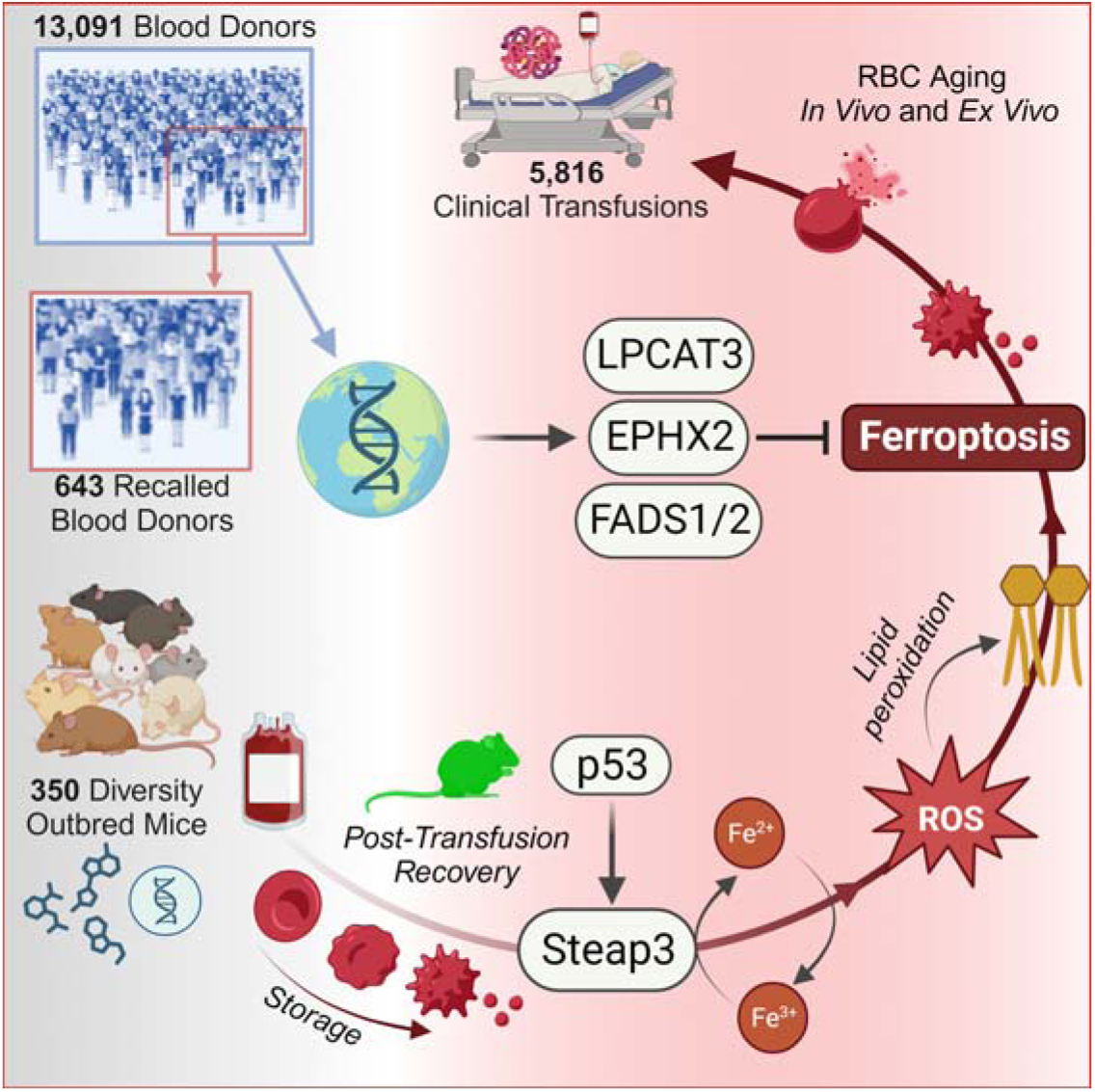

## Introduction

Garrod’s concept of ‘chemical individuality’, a founding principle of clinical biochemistry, fostered a century of research on the molecular origins of human diseases^1^. Fueled by discoveries of the genetic underpinnings of cross-individual heterogeneity in blood metabolism, clinical biochemistry and modern diagnostic medicine rely on the metabolic characterization of blood cells and biofluids as a window into an individual’s systemic metabolic health^2^. Red blood cells (RBCs) represent 99% of the circulating blood cells and 83% of the 30 trillion cells that compose an adult human^3^. RBCs are responsible for exchanging metabolites via >77 known active membrane transporters^4^ as they circulate through the body from the large arteries to narrowest capillaries. As such, RBC metabolism does indeed offer a unique window on systems health and dysregulation thereof as a function of genetic and non-genetic factors^5^.

Owing to their central role in gas transport, each RBC is loaded with ∼250-270 million copies of hemoglobin^6^. Hemoglobin is a tetrameric molecule whose conformation is tightly controlled by RBC metabolism via small molecule allosteric regulators that facilitate oxygen release through the stabilization of oxy/deoxyhemoglobin states^7^. Each hemoglobin subunit carries a heme group that coordinates a molecule of iron, resulting in up to ∼1 billion molecules of oxygen per RBC at full O_2_ saturation. Cumulatively, the total RBC “organ” accounts for up to 2.5 g of iron, which represents ∼66% of total bodily iron^8^. Every time an oxygen molecule binds to and then dissociates from hemoglobin, there is a 1:8 chance that it leaves hemoglobin with an extra electron, a process that generates reactive oxygen species (ROS) through ferrous-iron-promoted Fenton and Haber-Weiss chemistry^9^. Owing to the lack of nuclei and organelles, RBCs are incapable of replacing oxidatively damaged components through de novo protein synthesis. Therefore, in addition to their direct relevance to human health, RBCs offer an excellent, simplified eukaryotic cell model to investigate metabolic responses to oxidant damage – especially that arising from iron-dependent chemistry - in the absence of confounding transcriptional and translational buffers.

RBC transfusion is the most common, life-saving hospital medical procedure after vaccination (∼100 million units of RBCs every year), and blood storage is a practical necessity to meet transfusion demands. RBC blood unit’s storage life progresses to the limit of 42 days, the shelf-life indicated by Food and Drug Administration and European Council guidelines based on thresholds for in bag hemolysis and extravascular hemolysis, as determined via post-transfusion recovery (PTR) studies. During storage, RBCs accumulate ROS^10^ and oxidative damage to proteins and lipids, a progressively irreversible phenomenon,^11^ collectively referred to as the “storage lesion”. Such lesion promotes the vesiculation of damaged components, which in turn results in morphologically irregular^10^, smaller RBCs^12^ that are more susceptible to in-bag hemolysis, to intravascular hemolysis in the bloodstream or extravascular hemolysis via splenic and/or hepatic sequestration and erythrophagocytosis^12^. The latter process of extravascular hemolysis is also an essential regulator of the in vivo lifespan of RBCs^5^. To compensate for the inability to respond to oxidative stress and damage through de novo protein synthesis, RBCs are equipped with multiple antioxidant systems. These antioxidant pathways are regulated during blood storage through metabolic reprogramming: one paradigmatic example is the decrease in metabolic fluxes through glycolysis upon oxidation of glyceraldehyde 3-phosphate dehydrogenase, which favors the activation of the pentose phosphate pathway to generate NADPH, which in turn recycles the oxidized glutathione system back to its reduced form.^13^ While these mechanisms have been elucidated in small scale laboratory studies, larger scale clinical observations suggest that the onset, severity, and trajectory of the storage lesion is heterogeneous across blood donors, due to environmental factors (storage additives^14^, donor exposures),^15^ to biological (sex^16^, age^17^) or genetic factors (ethnicity^18^). Population studies in twins^19^ or large blood donors cohorts, such as the ∼13,000 donors enrolled in the Recipient Epidemiology and Donor Evaluation (REDS) Study^20^, are now shedding light on how genetics contribute to RBC storage quality. Current storage quality gold standards are, as defined by the U.S. Food and Drug Administration, are determined by measuring in-bag hemolysis (<1%) and PTR (>75%), i.e., the percentage of transfused RBCs that still circulate at 24h upon transfusion). Preliminarily, 27 genetic loci have been linked to in-bag hemolytic propensity^20^, and these polymorphic loci have also been linked to hemoglobin increments in recipients of donors carrying these alleles^21^. However, preliminary metabolite quantitative trait loci (mQTL) analysis of 250 blood donors suggests that this short list of polymorphisms barely scratches the surface of the molecular mechanisms that contribute to RBC metabolic adaptations to stress, including due to storage^22^. Genetic variants that impact RBC antioxidant capacity, such as glucose 6-phosphate dehydrogenase (G6PD) deficiency, confer susceptibility to hemolysis and post-transfusion performances of stored human RBCs^16,23^. Factors like donor ethnicity have been associated with lower propensity to hemolyze following osmotic insults (e.g., in donors of African descent)^18^, though a molecular rationale for this observation has not yet been identified.

Investigations of the genetic underpinnings of the metabolic and hemolytic heterogeneity in fresh and stored RBCs offer a window into the mechanisms that drive RBC responses to oxidant insults. Therefore, studies like the present one could inform strategies to design and test novel blood products for transfusions,^24^ or for novel interventions that benefit millions of patients suffering with hemolytic disorders (∼7% of humankind), from G6PD deficiency to sickle cell disease or pyruvate kinase deficiency^25^. To mechanistically explore these systems, our group has developed murine models of blood transfusion^26^, which we have leveraged to show how heterogeneous post-transfusion performances can be at least in part explained by cross-strain diversity in mice^27^. However, such studies have been hitherto limited to a handful of strains and relatively limited breeding strategies. The recently developed the Jackson Laboratory Diversity Outbred (J:DO) mouse model encompasses far greater levels of genetic diversity than traditional mouse crosses due to the intercrossing of eight highly divergent founder mouse strains, followed by extensive outbreeding^28^. This resource population can capture the impact of genetic heterogeneity in like fashion to population-based studies^29^, while still offering the benefits of murine experiments, including the standardization of relevant factors (e.g., age, sex, diet, exposures for RBC storage models) and strict control of environmental sources of variations^28,30^, enabling a next generation investigation into how genetic variation contributes to RBC metabolism and storage lesion heterogeneity.

## RESULTS

### Mapping Steap3 as a key genetic driver of RBC post-transfusion recovery in Jackson Laboratory Diversity Outbred (J:DO) mice

To map genetic variants that influence RBC storability in mice, 350 mice were selected from the 34^th^ generation of the ongoing J:DO breeding program (**Figure 1.A**). RBCs from these mice were stored for 7 days under conditions mimicking human RBC storage in the blood bank^26^. Stored RBCs were mixed with fresh mCherry tracer RBCs and then transfused into GFP+ mice. PTR was determined by establishing the test:tracer RBC ratio by flow cytometry on blood obtained 24 hours post-transfusion and normalizing to the pre-transfusion ratio (henceforth, post-transfusion recovery or PTR^31^ – **Figure 1.A**). Mice were genotyped on a ∼143,000 marker array^32^ and QTL analysis was carried out using PTR as a quantitative trait (**Figure 1.B**), which identified a strong QTL (peak LOD score = 36.3) on chromosome 1 (120.0 Mbp) nearby the encoded ferrireductase Steap3 (**Figure 1.C-D**).

**Figure 1.**
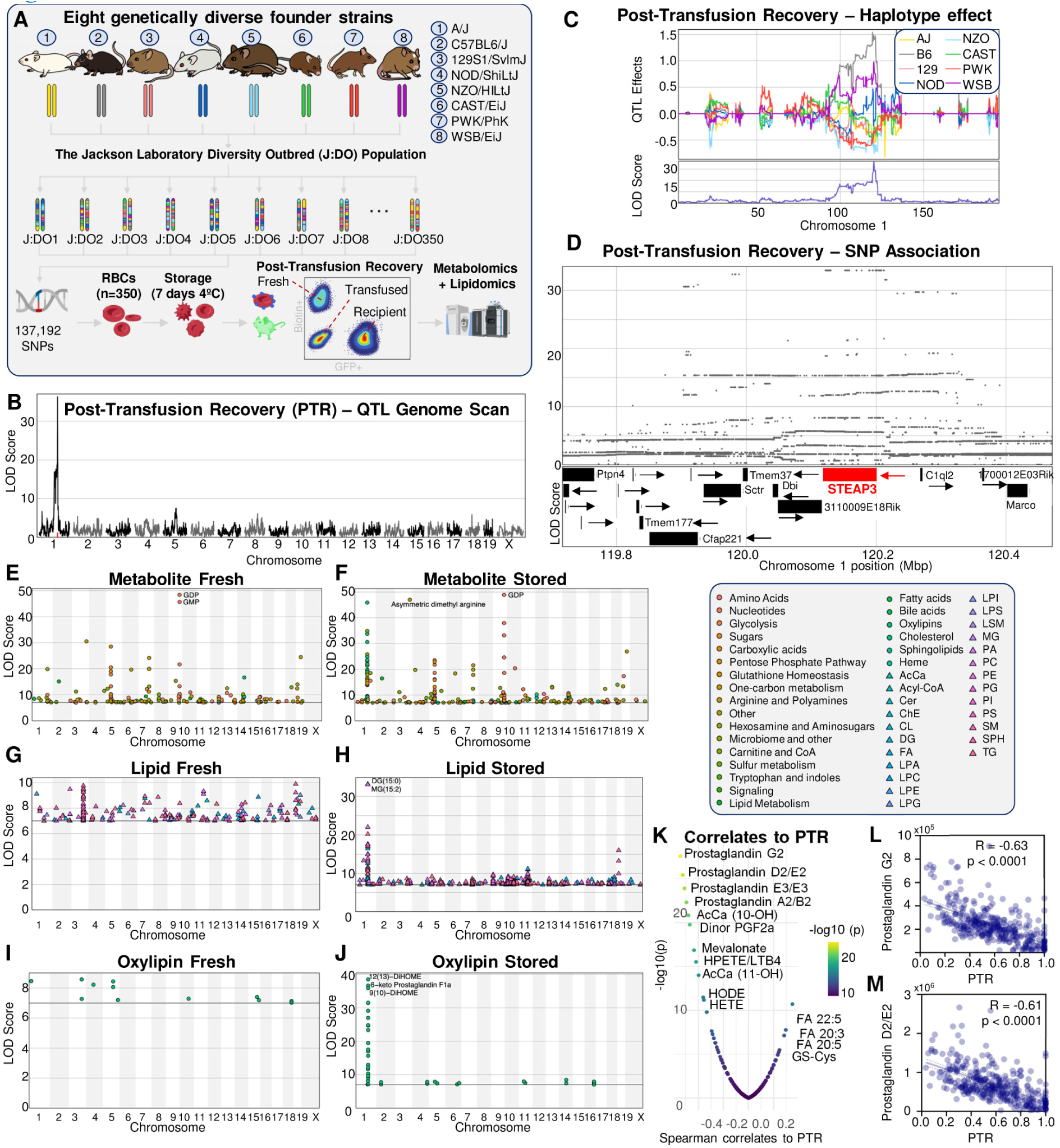
Genetic underpinnings of red blood cell (RBC) metabolic and storage quality heterogeneity in Jackson Laboratory Diversity Outbred (J:DO) mice. Diagram of genetically diverse mouse experiment (A). Eight founder strains were extensively intercrossed to produce highly recombinant outbred progeny. RBCs were collected and stored under conditions mimicking human RBC blood bank storage for 7 days, the end of shelf-life for pRBC in humans. RBCs were thus transfused into GFP+ mice to determine the percentage of transfused RBCs still circulating at 24h after transfusion (post-transfusion recovery [PTR]). Mice were genotyped and fresh and stored RBCs were analyzed via mass spec-based metabolomics and lipidomics. A quantitative trait locus (QTL) for PTR was mapped on chromosome 1 in the region encoding the ferrireductase Steap3, shown as the genome scan (B), haplotype effects at the QTL (C) and SNP associations in the QTL region (D). Combined Manhattan plots of peak associations for metabolites, lipids and oxylipins in fresh and stored RBCs reveal a hot spot region on chromosome 1 associated with changes in metabolite, lipid, and oxylipin levels specific to stored samples (E-J). Metabolites, lipids, and oxylipins with strong QTL are highlighted as labels. Molecular correlates to PTR identified lipid peroxidation products as strongly associated with PTR (K), including prostaglandin G2 and D2/E2 (isobars) (L-M).

### Genetic drivers of RBC metabolic heterogeneity in J:DO mice

Metabolomics and lipidomics analyses were carried out on a fresh and stored RBC sample from each of the 350 mice. Linear discriminant analysis (**Figure S1.A**), hierarchical clustering analysis of the top 50 metabolites by repeated measures ANOVA (**Figure S1.B**) and volcano plots (**Figure S1.C**) highlighted a significant effect of refrigerated storage on cell metabolism, in keeping with the prior literature^5^. Specific markers of the storage lesion included a significant decline in glycolysis (**Figure S1.D**), the only energy generating pathway in mature RBCs, and an increase in lipid peroxidation markers (**Figure S1.E**).

Genetic mapping analysis was then performed on the metabolomics and lipidomics data to identify QTL (here referred to as mQTL for metabolites and lQTL for lipids); two separate analyses were performed based on measurements in fresh and stored RBC samples (**Figure 1.E-J**). Based on a stringent threshold of LOD score > 8, we mapped 76 mQTL and 54 lQTL in fresh samples and 114 mQTL and 168 lQTL in stored samples. There were distinct regions of the genome where many QTL co-mapped, i.e., QTL hotspot regions (based on more than 5 QTL mapping within a sliding of window of 4Mbp). We observed mQTL hotspots on chromosomes 1, 5, 7, 9, 12, and 14 (plots of QTL density across the genome for fresh and stored metabolites, lipids and oxylipins are shown in **Figure S2.A-F**). For lQTL, we observed a fresh-specific hotspot on chromosome 3 and stored-specific hotspots on chromosomes 1, 7, 9, 10, 11, 13, and 19. Across metabolites, lipids, and oxylipins, the storage-specific QTL hotspots on chromosome 1 co-mapped with the PTR QTL.

We also imputed SNP and structural variant (SV)^33^ genotypes and performed association analysis (in contrast to the conventional haplotype-based approach commonly used as the initial QTL mapping in J:DO data) within QTL support regions. Notably, we identified a SV that was more strongly associated with thymidine levels than any SNPs, suggesting it may be the actual causal variant (**Figure S3.A**). The thymidine-associated SV was just upstream of the thymidine phosphorylase (Tymp) gene, an enzyme that catalyzes thymidine phosphorolysis. The QTL haplotype effects are highly consistent with an cis-eQTL for Tymp observed in liver tissue from another J:DO cohort^34^ (**Figure S3.F**), suggesting that J:DO mice with the B6 and PWK alleles at Tymp have low levels of Tymp protein, resulting in higher thymidine levels (**Figure S3.A-B**).

### Genetic variation at Steap3 is associated with post-transfusion recovery and lipid peroxidation

Oxylipins in stored samples (measured 24h post transfusion) are the top negative correlates of PTR (**Figure 1.K**). Specifically, prostaglandins (e.g., prostaglandin G2 and D2/E2 isobars - **Figure 1.L-M**) and other eicosanoid and octadecadienoic hydroxy- and hydroxy-peroxides (HETEs, HODEs and HPETEs, respectively) were amongst the top ten negative correlates to PTR. Conversely, very long chain poly- and highly unsaturated fatty acids (20:3; 20:5; 22:5) were positive correlates, suggesting that decreased storage-induced oxidation of these substrates is protective against a decline in post transfusion performance (**Figure 1.K**).

Looking across metabolites (including oxylipins) and lipids reveals an extensive QTL hotspot that covers the coding region for the ferrireductase Steap3 on chromosome 1 (**Figure 2.A-B**). We used the QTL results to define a co-mapping network view of the gene-metabolite pairs, which illustrates the centrality of the Steap3 QTL hotspot (**Figure 2.C**). An mQTL density plot highlights the extent of metabolites, lipids, and oxylipins regulated by Steap3 in stored samples (**Figure 2.D-F**). The haplotype effects at this QTL region are complex with alleles from B6 and WSB being strongly distinct from the others, shown here for hydroxyoctadecadienoic acid (HODEs – combined isomers unresolved with the high-throughput metabolomics method but further resolved by the independent oxylipin analysis – **Figure 2.G**). Steap3 is located centrally within the QTL region and SNP variants with alleles shared by B6 and WSB best capture the haplotype association (**Figure 2.H**). The genetic effects are highly consistent and specific to stored samples within this QTL hotspot, which is emphasized by how the relationships between metabolites, lipids, oxylipins, and PTR resolves into two correlated and anti-correlated groups and suggests they are driven by the same genetic variation at Steap3 (**Figure 2.I-K**). Given the technical challenge of separating isobaric isomers for oxylipins, which ranked as the top correlates of PTR, we used an ad hoc method^35^ to orthogonally separate and measure oxylipins, which confirmed a strong signal for oxylipin accumulation in stored RBCs as a function of Steap3 SNPs (**Figure 2.F**).

**Figure 2.**
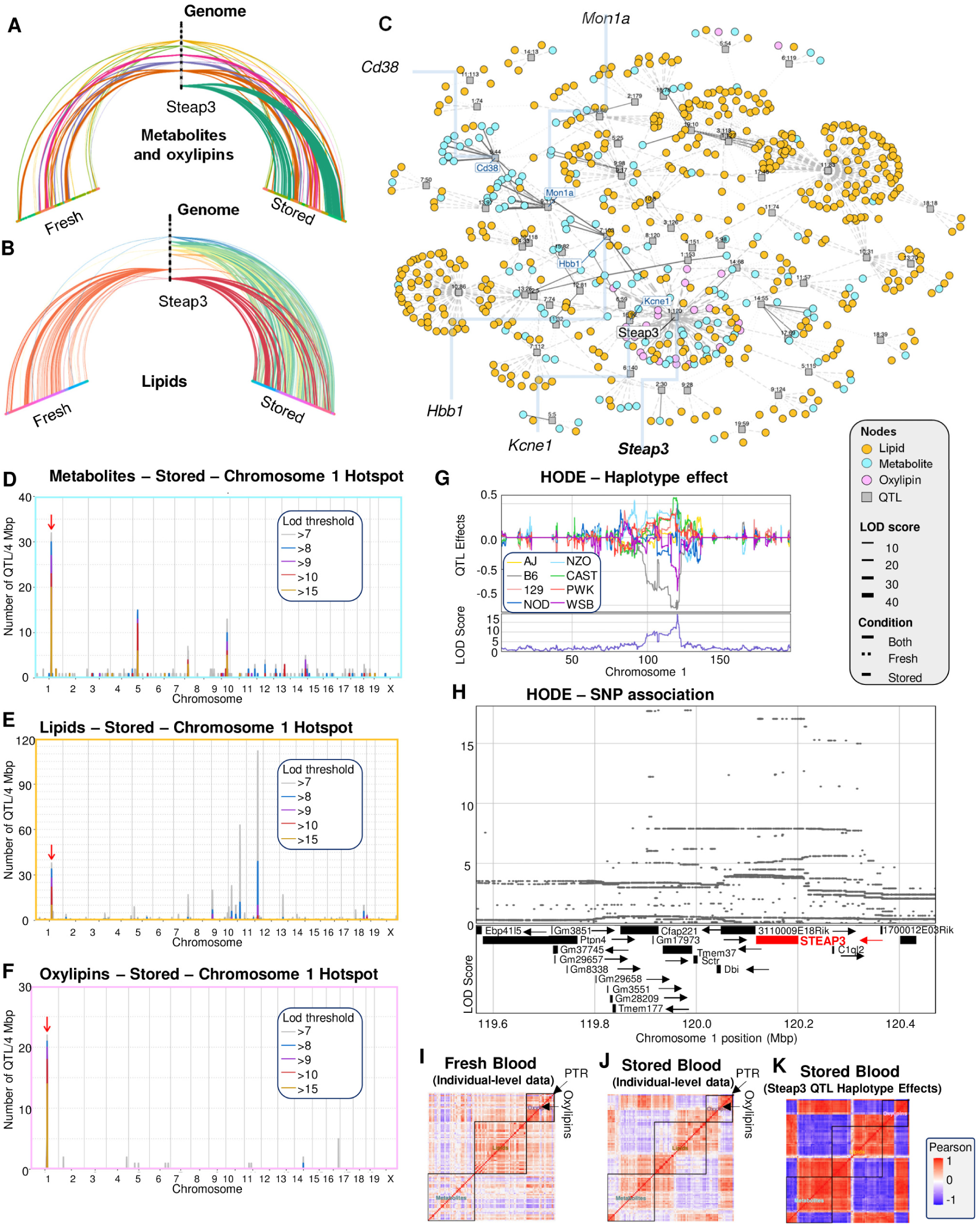
Chromosome 1 QTL hotspot reveals Steap3 as a key regulator of metabolites, oxylipins and lipids in stored RBCs. QTL for metabolites, oyxlipins and lipids revealed co-mapping hotspots, including the STEAP3 locus on chromosome 1 specific to stored RBCs, depicted as hive plots (**A**-**B**), a network plot (**C**), and QTL density plots (**D**-**F**). 3D model of the membrane proximal oxidoreductase domain of human Steap3 (bound to NADP) is overlayed with the chromosome 1 hotspot. Other candidate gene drivers of hotspots are included as labels. Hydroxyoctadecadienoic acid (HODE) is an example of a metabolite that maps the Steap3 QTL (chromosome 1 scan and haplotype effects – **G** and SNP associations – **H**). The correlation structure among metabolites, lipids, oxylipins, and PTR reflects the Steap3 QTL, with fresh samples being uncoordinated across data types (**I**), but highly coordinated in stored samples, at both the individual-level data (**J**) and the QTL haplotype effects (**K**).

### A metabolomics and lipidomics QTL resource webtool

While not the main focus here, these data and QTL results represent a powerful resource for surveying the effects of genetic diversity on metabolite and lipid levels. We have developed a publicly available online portal using our QTLViewer webtool^36^ for interactive exploration of QTL results and distribution of the processed data, accessible at: https://churchilllab.jax.org/qtlviewer/Zimring/RBC.

Functionality includes plotting and exporting results from PTR, metabolite and lipid phenotypes, including the ability to inspect how covariates of interest (e.g., sex) correlate with the traits, QTL scans, SNP association, and haplotype effect scans. We also identified strong candidate genes at some of the other QTL hotspots, which are visualized as hive plots and network graphs in **Figure 2.A-C**, and more extensive follow-up in **Figures S4-S12.** Other than the hotspot on chromosome 1, we identified additional metabolite-gene associations including (i) glycolysis ^37^ and pentose phosphate pathway metabolites and the region on chromosome 5 coding for CD38/BST1 – **Figure S4**); (ii) antioxidant metabolites and a region on chromosome 7 mapping on polymorphic hemoglobin beta (Hbb1 and Hbb2 – **Figure S5**); (iii) purine nucleosides and a region on chromosome 9 mapping on MON1 homolog A (Mon1a – **Figure S6**); and (iv) lysine metabolites and a region on chromosome 14 (**Figure S7A-E**).

Similarly, lipid-gene associations were identified for lipids extracted from fresh RBCs and a region mapping on chromosome 3 (**Figure S7.F-I**), or lipids extracted from stored RBCs and regions on (i) free fatty acids (18, 20 and 22 C aliphatic chain length); (ii) oxo-diacyl- and oxo-triacyl-glycerols and chromosome 7 (**Figure S8.A-D**), (iii) phospholipids and chromosome 9 and 10 (**Figure 8.E-H** and **Figure S9.A-D**, respectively); (iv) sphingolipids, triacylglycerols and additional hotspots on chromosome 10 (**Figure S9.E-H** and **Figure S10.A-D**); lysophospholipids and chromosome 11 (**Figure S10.E-H**); (vi) triacylglycerols and chromosome 13 (**Figure S11.A-D**); (vii) oxo-lysphophospholipids and chromosome 18 (**Figure S11.E-H**). The depth of these secondary findings highlights the value of the J:DO population as a resource for metabolomics and lipidomics.

### Genetic manipulation of STEAP3 regulates lipid peroxidation in stored murine RBCs

Our findings from the J:DO population on PTR independently validated previous work from our groups^31^ that used an F2 cross-generation between just two strains of mice: C57BL/6J mice, previously identified as good storers^27^, and FVB/J mice, previously identified as poor storers^27^. This analysis identified the Steap3 locus at lower resolution (3Mb in Howie et al.^27^, here further narrowed down to 2.85Mb after additional 6 generations of backcrossing to C57BL/6J) – as confirmed by strain-specific ferrozine assays - that impacts lipid peroxidation in stored murine RBCs (**Figure S13.A-D**). Given that the J:DO population included C57BL/6J as founder but not FVB/J, this study allowed us to characterize the effects of new Steap3 alleles and potentially map additional genetic loci for PTR. Though we did not map new additional QTL for PTR, the haplotype effects at Steap3 are complex. Bayesian modeling of the Steap3 QTL for PTR founder evidence for more than two functional alleles in the J:DO^38^, suggesting multiple genetic variants at Steap3 affect PTR and downstream metabolites/lipids (**Figure S12**).

Independent, unsupervised validation of the Steap3 finding as a lynchpin of poor PTR via lipid peroxidation with two separate murine studies prompted us to follow up on the observational results with mechanistic interventions. To this end, first we knocked in the hypermorphic STEAP3 from FVB/J mice into C57BL/6J mice, which increased lipid peroxidation (Volcano plots highlight oxylipins in red – **Figure S13.E**) and decreased PTR (∼25%) in good storing mice (**Figure S13.F**). We then investigated STEAP3 knockout (KO) mice, which also had a significant impact on RBC metabolism upon storage (**Figure S13.G**). Specifically, both Steap3 KO and hypermorphic gain of function (GOF) knock in (KI) resulted in decreases in PTR in mice (∼45%), which was still two-fold higher in Steap3 KO compared to KI mice (**Figure S13.H**). However, only GOF KI mice but not KO show elevated levels of oxylipins in end of storage RBCs (**Figure S13.I**), demonstrating that Steap3 presence/activity in mature RBCs is required to promote lipid peroxidation and quality decline in stored RBCs.

### Oxylipins, iron and STEAP3 are associated with hemolysis in 13,000 REDS RBC Omics blood donors

The roles of Steap3 in lipid peroxidation and decline in storage and post-transfusion quality, as identified in murine models, have not yet been validated in humans. To address this limitation, we first determined the metabolic underpinnings of hemolysis measurements in 13,091 donors from the REDS RBC Omics studies (**Figure 3.A**). We identified oxylipins (particularly, HETEs and HODEs) as top correlates of end of storage hemolytic propensity in human packed RBCs (**Figure 3.A**). To expand on these findings, we then reached out to the original (index) cohort and obtained a second independent unit of packed RBCs from 643 volunteers who had shown extreme hemolytic propensity (5^th^ vs 95^th^ percentile) during the index screening phase (**Figure 3.B**). We determined the metabolic correlates to vesiculation rates – a hallmark of increased oxidative damage to the RBCs ^11^ and a mechanism driving decrease in RBC size, a predictor of post-transfusion extravascular hemolysis via splenic sequestration^12^ - in the 643 units at storage day 10, 23 and 42 (representing 1,929 total samples). This process confirmed HETEs and HODEs as the top markers of RBC vesiculation (**Figure 3.B**).

**Figure 3.**
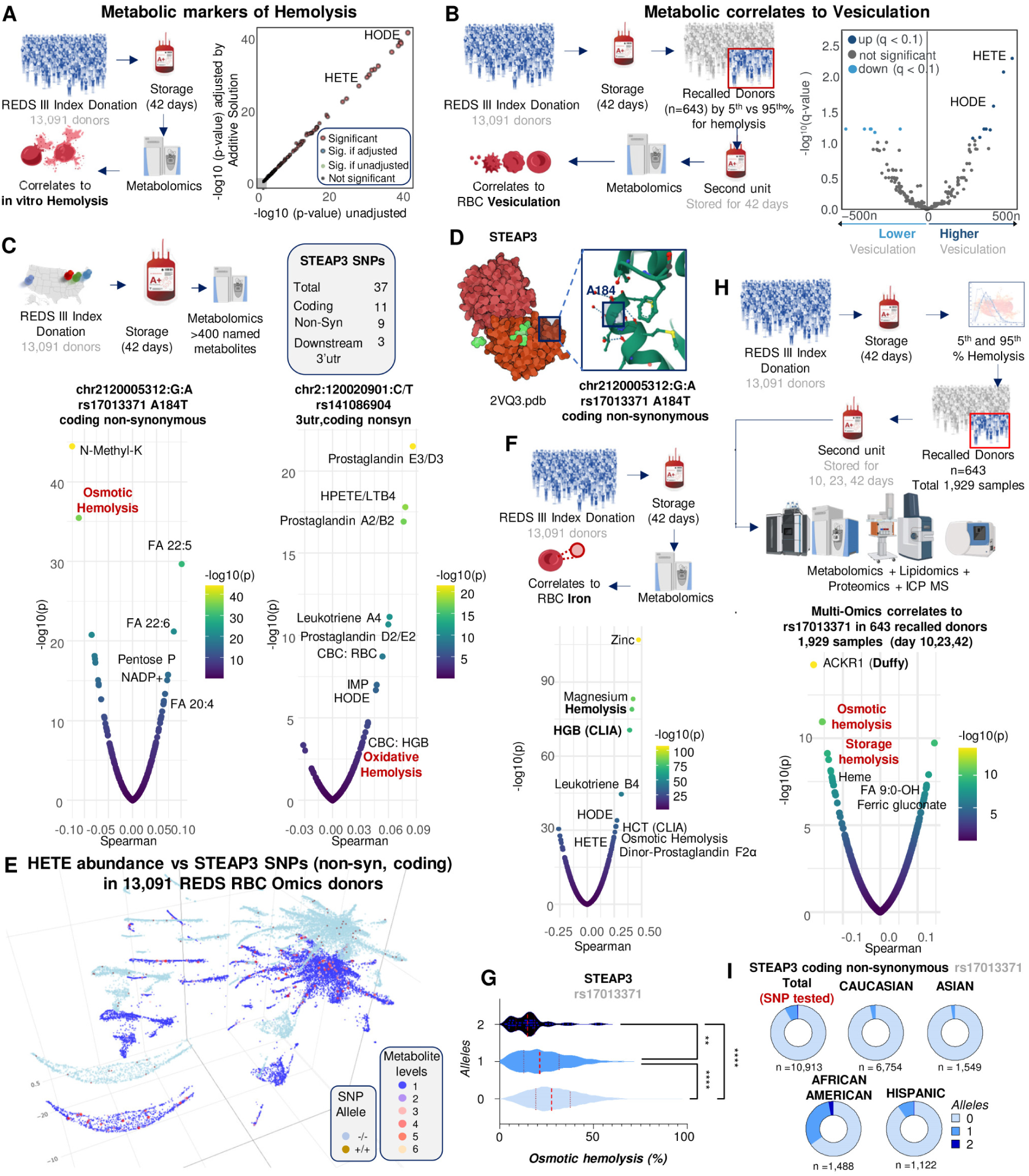
Oxylipins, iron and STEAP3 are associated with hemolysis in 13,000 REDS RBC Omics blood donors. Hemolysis measurements in 13,091 donors from the REDS RBC Omics studies identified oxylipins (especially, HETEs and HODEs) as top predictors of end of storage hemolytic propensity (**A**). A second independent unit from 643 of the index donors who were contacted because of their extreme hemolytic propensity (5^th^ vs 95^th^ percentile). Determination of vesiculation rates in the 643 units at storage day 10, 23 and 42 identified HETEs and HODEs as top markers of RBC vesiculation (**B**). In **C**, 37 SNPs were monitored for STEAP3 in the REDS RBC Omics study. Common non-synonymous coding SNPs correlated with osmotic and oxidative hemolytic propensity and oxylipin levels (**C**). Some of these SNPs were predicted to impact the ferrireductase function of the enzyme (**D**). 3D uMAP representation of HETE levels and all non-synonymous STEAP3 SNPs in homozygosity are shown in **E**. In **F**, iron measurements via ICP-MS in the REDS RBC Omics recalled donor cohort showed significant association with hemolysis and oxylipins. Hemolysis was significantly lower in donors carrying two alleles of rs17013371, the most common non-synonymous STEAP3 coding SNP (**G**). Multi-omics correlates in the same recalled donor cohort confirmed such association for the rs17013371 (**H**), a SNP that is over-represented in donors of African American descent.

After confirming that lipid peroxidation is a marker of hemolysis and vesiculation in stored human RBCs, we set out to determine whether there are associated genetic variants at Steap3 in human blood donors. First, we identified and evaluated 37 SNPs annotated to Steap3 in the 13,091 REDS RBC Omics index donors (**Figure 3.C**). A full list and summary of allele frequency is provided in **Figure S14**. A majority of the SNPs were intronic (23). More interestingly, 11 of the SNPS were coding, of which 2 were synonymous and 9 were non-synonymous. The final 3 were in the downstream 3^’^-UTR. Steap3 variants were associated with hemolysis in stored human RBCs; for example, rs17013371 was associated with RBC susceptibility to hemolysis upon osmotic insults (**Figure 3.C**). Similarly, rs141086904 (**Figure 3.C**) and rs147820529 (**Figure S14.B**) were associated with multiple prostaglandins and oxylipins (e.g., HODEs), as well as an increased susceptibility to hemolysis following oxidant insults. These common variants are located in regions neighboring the active site of the Steap3 enzyme (e.g., rs17013371 results in the A184T substitution), which is predicted to impact enzyme kinetics (**Figure 3.D**). To visualize the association between Steap3 SNPs and lipid peroxidation, we generated a figure by superimposing two identical 2D uMAPs based on metabolomics results for the whole REDS population: the bottom map was color-coded based on HETE levels and the top one based on genotypes, specifically homozygosity for any of the non-synonymous STEAP3 SNPs (**Figure 3.E**).

Steap3 is a ferrireductase required for efficient transferrin-dependent iron uptake in erythroid cells^39^ and thus regulates intracellular iron content in mature RBCs. Based on this mechanism, we predict that elevated iron content, especially in its ferrous state as a result of Steap3 activity, would promote Fenton and Haber-Weiss chemistry in iron-loaded erythrocytes, which in turn would favor lipid peroxidation via ferroptosis^40^, also referred to as “death by lipid peroxidation”^41^. To evaluate this, we measured RBC iron levels via ICP-MS in the REDS RBC Omics recalled donor cohort (n=643 – **Figure 3.F**). Consistent with our hypothesis, correlation of omics data to RBCs iron levels revealed a significant association with hemolysis and oxylipins (**Figure 3.F**). Carrying one or two alleles of non-synonymous coding SNPs for Steap3 (e.g., rs17013371 – 2 alleles) was negatively associated with osmotic and storage hemolysis in both the index and recalled donor cohort (**Figure 3.G-H**). In the latter group, combined multi-omics analyses (including proteomics, lipidomics, oxylipins, and trace element analysis) identified an association between this SNP and RBC heme and iron content (ferric gluconate - **Figure 3.H**), as well as oxononanoic acid (FA 9:0-OH), a breakdown product of FA 18:2-derived oxylipins^42^ (e.g., HODEs). Interestingly, the Steap3 rs17013371 SNP was also negatively associated with elevated levels of the Duffy blood group protein atypical chemokine receptor 1 (ACKR1 – **Figure 3.H**). Of note, rs17013371 alleles were significantly more prevalent in donors of African American descent (**Figure 3.I**).

### Genetic factors contribute to heterogeneous lipid peroxidation in 13,091 human RBCs after storage for 42 days

To determine the genetic underpinnings of lipid peroxidation at end of human RBC storage and expand on our results in mice, we performed genome-wide association analysis for oxylipins. Specifically, we performed targeted measurement of oxylipins in 13,091 end of storage packed RBC units from the REDS RBC Omics index donor cohort and performed mQTL analyses based on 879,000 SNPs from a precision transfusion medicine array^43^ (**Figure 4.A**). Representative Manhattan and LocusZoom plots are shown for HETE, HPETE, Leukotriene A4, prostaglandin A2, dinor-prostaglandin F2α, prostaglandin D2/E2 and G2 (**Figure 4.A-K**). The combination of oxylipin-gene hits elucidates a pathway of lipid detoxification fueled by L-carnitine,^44^ NADPH and glutathione-dependent systems (**Figure 4.I**), a map that substantially overlaps with literature in the field of ferroptosis^40^.

**Figure 4.**
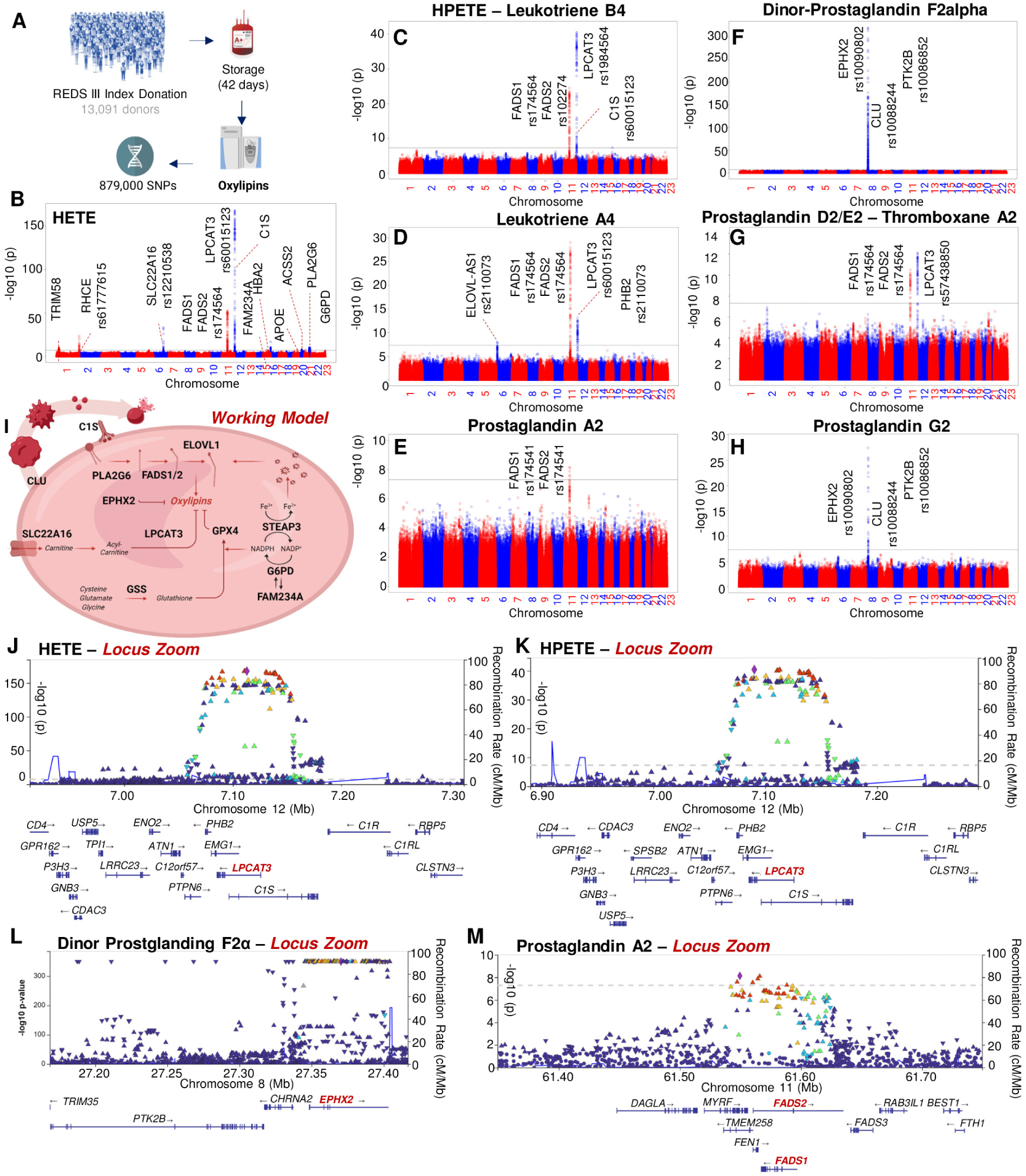
Genetic factors contributing to heterogeneous lipid peroxidation in 13,091 human RBCs after storage for 42 days. Genome-wide association studies (GWAS) were performed for lipid peroxidation products in the REDS RBC Omics index donor cohort (n=13,091) against 870,000 SNPs from a precision transfusion medicine array (**A**). From **B-H**, Manhattan plots are shown for HETE, HPETE, Leukotriene A4, prostaglandin A2, dinor-prostaglandin F2alpha, prostaglandin D2/E2 and G2. In **I**, a summary model of the pathway that emerges as a regulator of lipid peroxidation in stored human RBCs. In **J-M,** representative locus zoom plots for selected top hits by significance (-log10 p-values, y axes).

Of all oxylipins, HETEs (combined isomers – **Figure 4.B**) showed the highest number of hits, including polymorphisms in the regions coding for lysophosphatidylcholine acetyl-transferase 3 (LPCAT3 – rs60015123 p=1.54E-168); fatty acid desaturases 1 and 2 (FADS1 or 2; rs174564 p=1.38E-53); the carnitine transporter SLC22A16 (rs12210538; p= 3.03E-34); glutathione synthetase (GSS; rs6087652 p=1.88E-08); Phospholipase A2 Group VI (PLA2G6; rs3827354 p=4.82E-11) among other hits, also including the rate-limiting enzyme of NADPH-generating pentose phosphate pathway, glucose 6-phosphate dehydrogenase (G6PD, rs1050828, c.202G>A; p.Val68Met, known as the “common African variant”) and FAM234A (associated with G6PD deficiency in population studies^45^). Most of these hits were shared with other oxylipins, including HPETE, prostaglandin A2 and D2/E2, leukotriene A4 (**Figure 4**), HODEs (**Figure S15**). Additionally, epoxide hydrolase 2 (EPHX2) emerged as the most significant hit for dinor-prostaglandin F2α (rs10090802; p<e-308 – **Figure 4.F**) and prostaglandin G2 (**Figure 4.H**).

The strongest for oxylipins and genetic variations in LPCAT3, FADS1/2 and EPHX2 were confirmed by LDA in the index donor cohort (**Figure 5.A-C**) and validated in the recalled donor cohort (**Figure 5.D-F**), where top positive and negative features associated to allele frequency recapitulated all the top hits from the analysis of the Steap3 rs17013371 and rs141086904, though with higher significance. Similar to Steap3 SNPs, allele frequencies for LPCAT3 rs60015123 (nonsense mutation associated with increased protein decay), FADS1/2 rs174564 (upstream variant) and EPHX2 rs10090802 (regulatory region variant) were significantly associated with lower (EPHX2 and LPCAT3) and higher (FADS1/2) hemolytic propensity upon osmotic insult (**Figure 5.G-I**). Also consistent with Steap3, breakdown of allele frequency by ethnicity showed a significant enrichment in blood donors of African descent for LPCAT3 and EPHX2, while FADS1/2 was underrepresented (**Figure 5.J**). Again, consistent with multi-omics correlates of Steap3 in the recalled donor population, SNPs in all genes associated with oxylipins in stored human RBCs are also associated with the Duffy blood group antigen protein ACKR1 **– Figure S16.A**).

**Figure 5.**
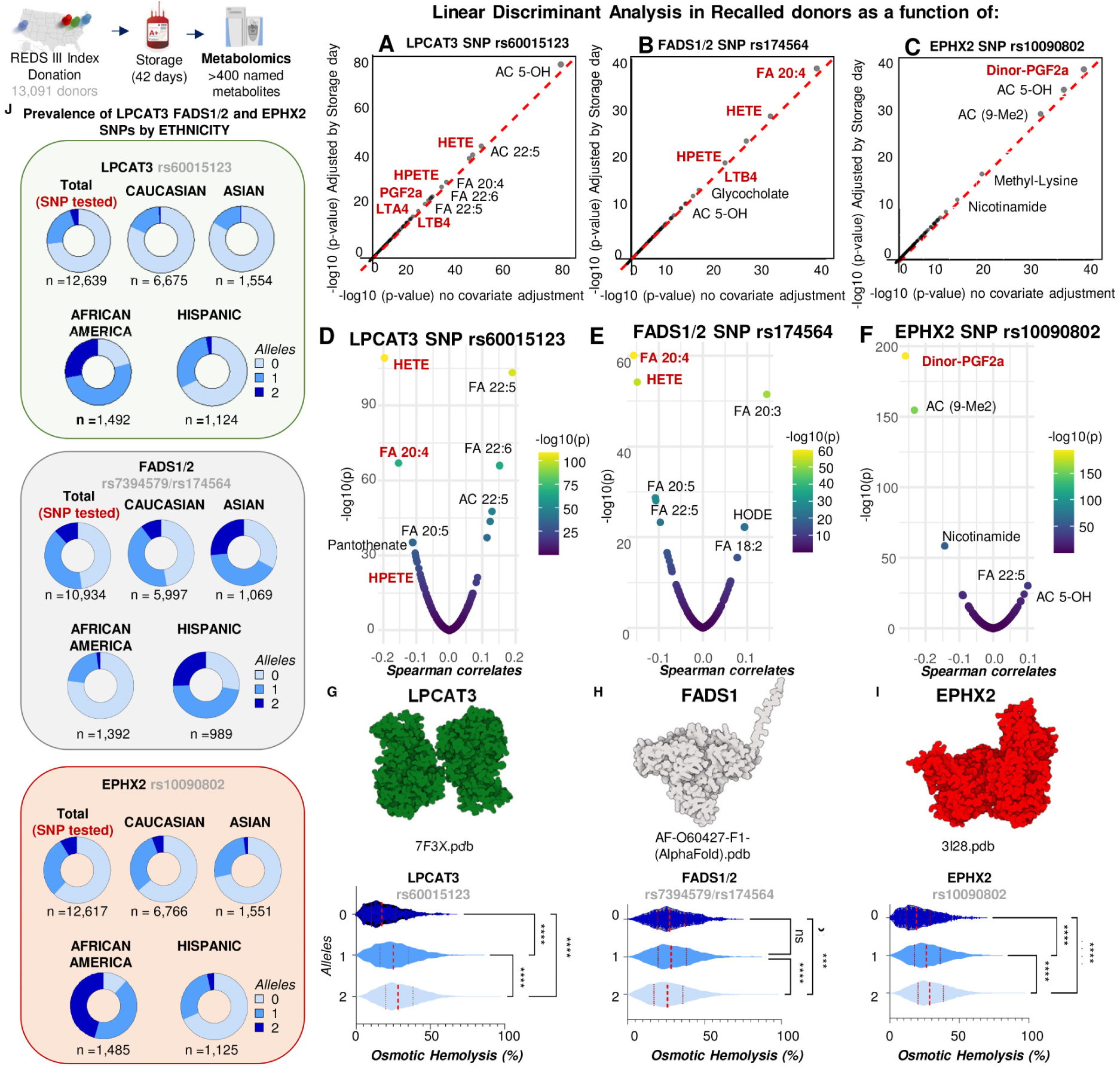
Genetic polymorphisms in LPCAT3, FADS1/2 and EHPHX2. Analyses in the recalled donor cohort confirmed findings from the index donor studies showing a strong association between SNPs in LPCAT3, FADS1/2 and EPHX2 and lipid peroxidation products via linear discriminant analysis (unadjusted or adjusted by storage day - **A-C**) and Spearman correlation (**D-F**). Allele dosage was associated with a decrease in hemolytic propensity (upon osmotic insult) in the larger index donor cohort (**G-I**). Breakdown of allele frequency by ethnicity showed a strong enrichment in blood donors of African descent for LPCAT3 and EPHX2, while FADS1/2 was underrepresented in this ethnic group (**J**).

### TP53 is polymorphic in the healthy blood donor population and associates with genetic regulators of lipid peroxidation and hemolysis

While Steap3 did not directly emerge as a hit from the mQTL analysis of oxylipins in humans, significant loci do point to a role for enzymes involved in ferroptosis protection as critical regulators of RBC lipid peroxidation in stored human RBCs (**Figure 4.I**). Steap3 is a transcriptional target of tumor suppressor protein TP53, which is also known as tumor suppressor-activated pathway 6 or TSAP6^46^. The role of TP53 in the regulation of ferroptosis has been extensively investigated^40,47^. Both p53 and STEAP3 are critical to erythropoiesis^48–50^ and polymorphic in humans: the prevalence of p53 germline mutations is estimated ∼1:2,000 births^51^ and somatic mutations accumulate with age^51–54^. While the role of p53 has been extensively studied in normal and malignant hematopoiesis^55^, little is known whether p53 is polymorphic in healthy human blood donor volunteers and whether such polymorphisms affect mature RBC metabolism and hemolysis.

To address this gap, we evaluated over 50 common TP53 SNPs in the 13,091 REDS RBC Omics blood donors (the top 20 by allele frequency are plotted in **Figure 6.A**). The list includes common SNPs associated with increased likelihood to develop cancers, such as rs1042522 (associated with the P72R mutation), rs150200764 (also known as the hereditary cancer predisposing syndrome, benign), rs1800371 and rs1800372 (H178R mutation in the DNA-binding site of TP53 – **Figure 6.A**). Notably, a non-coding SNP, rs8064946, was found to associate with osmotic hemolysis, as well as with the most significant hits from the mQTL analysis for human and murine oxylipins, including EPHX2, LPCAT3, G6PD, Steap3, GPX4 and SLC22A16, in order of significance from LDA analyses in REDS recalled donors (both unadjusted or adjusted by storage day – **Figure 6.B**). In both index and recalled donors (**Figure 6.C-D**), rs8064946 was associated with markers of osmotic fragility (FA 22:5 and FA:6; kynurenine) and G6PD-dependent pentose phosphate pathway cofactors and end-products (pentose phosphate – combined isomers, NADP+). Allele frequency for the rs8064946 TP53 SNP was positively associated with SNPs for EPHX2, LPCAT3, and negatively associated with G6PD, SLC22A16 and the Duffy blood group protein ACKR1 (**Figure 6.D**). Allele frequency as a function of ethnic breakdown revealed a lower frequency of the potentially pathogenic mutations rs1042522 and rs105200764 and higher frequency of the non-coding rs8064946 in donors of African descent (**Figure 6.E-H**). All these variants were associated with lower osmotic fragility (**Figure 6.I**). Relevant to the potential role of precision transfusion medicine array-based characterization of blood donor populations, the genotypes of the pathogenic TP53 SNPs, rs1800371 and rs1800372 (H178R and R248Q mutations in the DNA-binding site of TP53 – **Figure 6.I** and **Figure S16.B**), only three blood donors were observed that were homozygous, while heterozygous genotypes were far more frequent. This potentially pathogenic TP53 SNP is not only associated with the functional R248Q mutation, but also observed here to associate with elevated susceptibility to hemolysis following osmotic insults (**Figure 6.I**).

**Figure 6.**
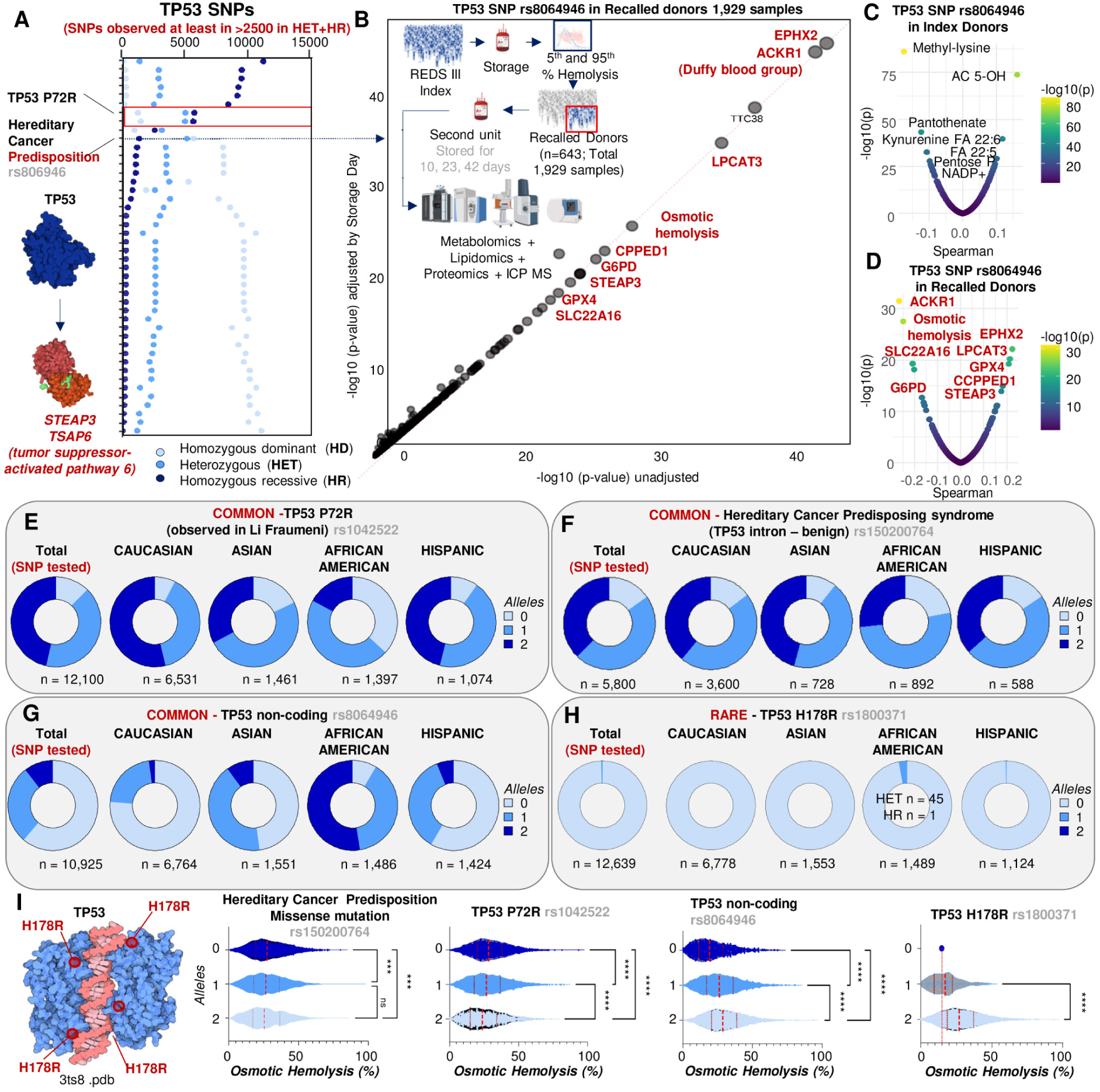
TP53 is polymorphic in the healthy blood donor population and associates with hemolysis. Over 50 common TP53 SNPs were monitored in the 13,091 REDS RBC Omics blood donors (top 20 plotted **A** as a function of allele frequency), including common SNPs associated with increased likelihood to develop cancers. One of these SNPs was identified as the top TP53 SNP associated with hemolytic propensity, a SNP whose allele frequency significantly correlated with that of the alleles for all the genes contributing to lipid peroxidation from the GWAS analysis (**B**). In index (n=13,091 - **C**) and recalled donors (validation cohort - n=643; **D**), the TP53 rs8064946 was associated with markers of osmotic fragility, oxylipins and poly-unsaturated fatty acids, as well as to allele frequency and protein levels for all lipid peroxidation-associated genes, when detected via proteomics (**D**). In **E-H**, allele frequency as a function of ethnic breakdown shows a lower frequency of the potentially pathogenic mutations rs1042522 and rs105200764 and higher frequency of the non-coding rs8064946 in donors of African descent. All these variants were associated with lower osmotic fragility (**I**).

### Genetic variants that regulate oxylipins are also associated with in vivo hemolysis of critically ill individuals receiving non-autologous transfusion

To investigate the potential clinical relevance of our findings, we accessed the “vein-to-vein” database of the REDS RBC Omics program^17^ and linked blood donor genotypes and end of storage oxylipin levels to the hemoglobin increments in thousands of single unit transfusion recipients of units donated by REDS donors. First, we observed that transfusion of units from donors who carried two copies of the rs8064946 TP53 SNPs was associated with significantly lower hemoglobin increments (p < 0.001), particularly when the transfused unit was older than 5 weeks (n=4,636 - **Figure 7.A-B**). Similar observations were made when focusing on transfusion of units from donors carrying two copies of the LPCAT3 rs60015123 allele, with significantly lower Hgb increments both immediately after transfusion and at 24h when transfused units were older than 3 weeks (n=4,473 transfusion events - **Figure 7.C-D**); this phenomenon was exacerbated by prolonged storage (i.e., 36-42 days) of transfused units for LPCAT3 rs60015123 homozygous donors (**Figure 7.E**). Finally, we show significant decreases in hemoglobin increments in recipients of units from donors with elevated end of storage oxylipin levels. Specifically, we saw that elevated day 42 HETEs were associated with significantly lower hemoglobin increments upon heterologous transfusion to critically ill recipients requiring transfusion (n=1,241 - **Figure 7.F**).

**Figure 7.**
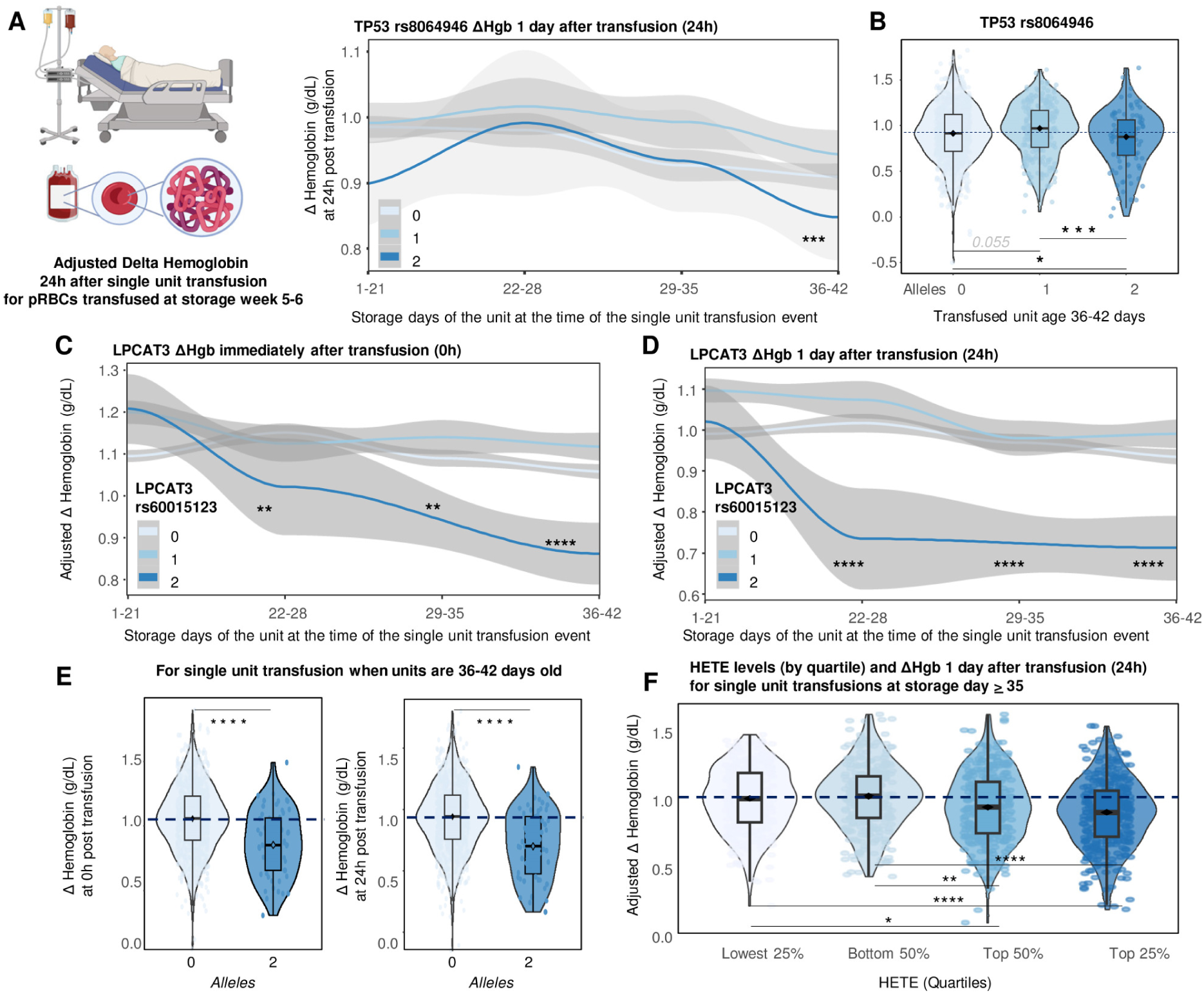
Oxylipins and genetic polymorphisms that regulate them are associated with in vivo hemolysis critically ill individuals receiving non-autologous transfusion. By accessing the “vein-to-vein” database of the REDS RBC Omics program, we linked blood donor TP53 rs8064946 status (homozygous dominant = HD; heterozygous = HET; homozygous recessive = HR) to adjusted hemoglobin increments in clinical recipients of single pRBC unit transfusions of different storage ages (**A**) or at storage day 36-42 (**B**). Similar analyses were performed for LPCAT3 rs60015123, focusing on Hgb increments immediately after transfusion (**C**) or at 24h (**D**) at any storage age, or for transfused units aged 36-42 days (**E**). Similarly, we linked end of storage oxylipin levels – e.g., hydroxyeicosatetraenoic acid, HETE – by quartiles to hemoglobin increments upon heterologous transfusion of products from the same donors to critically ill recipients requiring transfusion (**F**).

## DISCUSSION

In recent years, high-throughput technologies have enabled mQTL studies that delve into the genetic underpinnings of metabolic heterogeneity of plasma and serum^1,56^. In this study, we generated a novel resource that encompasses over 400 QTL from fresh and stored RBC samples from 350 genetically diverse mice, at baseline and following oxidant stress upon storage under blood bank conditions. The publicly available portal associated with the present data will enable ease access and interrogation of our findings for future studies focusing on gene-metabolite interactions. The focus of the present study on RBC metabolism stems from the opportunity to leverage a simplified mammalian cell system, yet one that mirrors systems metabolism and regulates it by participating in critical gas homeostasis. Indeed, the circulation and proper function of RBCs are required for the health of essentially all tissues^57^. RBCs deliver O_2_, scavenge CO_2_, engage in additional complex gas exchange and participate in regulating vascular tone (e.g. nitric oxide^58,59^ and ATP release^60^) and systems metabolism^8^. These processes are regulated metabolically, owing to RBC inability to respond to energy and redox challenges via de novo protein synthesis and/or epigenetic regulation of gene expression^5^. Thus, understanding the genetic underpinnings of RBC metabolism at baseline or upon oxidant challenge is of general significance to health and disease, since oxidant stress to RBCs is aggravated when oxygen demand is high, for example in response to exercise^61^, and numerous diseases have RBC dysfunction and hemolysis as part of their pathology^62^.

Altered RBC antioxidant capacity triggers intra-^16^ or extra-vascular hemolysis^12,63^, resulting in excess circulating heme and iron^64^, a phenomenon that underlies the etiology of many diseases in which oxidant stress plays a central role. Increased intra- or extravascular hemolysis alter iron and heme metabolism, which – combined with impaired O_2_ kinetics as a result of RBC dysfunction – promote damage of high-oxygen consuming organs kidneys^65,66^, heart^67–69^, lung^70–72^ and brain^73,74^. Strategies that mitigate oxidant stress to RBCs hold immediate critical biomedical implications, especially in the field of transfusion medicine, with RBC transfusion saving the lives of 4-5 million Americans every year. Dysregulated RBC metabolism under refrigerated storage conditions promotes increases in hemoglobin oxygen saturation and fuels the generation of reactive oxygen species, triggering the so-called storage lesion^11^. As such, understanding the genetic underpinnings of the metabolic heterogeneity of stored murine and human RBCs is essential to improving storage strategies and transfusion efficacy, but also a translationally relevant model of RBC responses to oxidant stress as they age in vitro. With an average in vivo lifespan of 120 days, every day ∼0.2 trillion RBCs are removed from the bloodstream, while as many are generated from hematopoietic stem cell precursors. Erythropoiesis relies on the uptake of circulating iron in the ferric (oxidized) state, and its intracellular reduction to the ferrous state, a process catalyzed by the ferrireductase STEAP3^39^, a transcriptional target of tumor protein TP53^75^.

Genetic manipulation of STEAP3 has been associated with anemia in mice^48^, but common polymorphisms in humans of Asian descent have not been associated with phenotypic changes^52^. In mice and humans, STEAP3 plays a central role in the regulation of iron metabolism via endosome trafficking^76^ and exocytic transport via vesiculation^46^. Recently, we had performed genetic studies on murine models of in vitro aging of RBCs and documented that – of 20 open reading frames mapping on a 3 Mb region on chromosome 1 – the hypermorphic STEAP3 represented the most likely candidate explaining poor blood storage quality in certain mouse strains (e.g., FVB) as compared to other strains with good storage (e.g., C57BL/6J)^77^. The aim of the present study was to identify new genetic factors that contribute to extravascular hemolysis of stored, transfused RBCs. To do so, we leveraged a larger cohort (n=350) of different mouse strains (not including FVB/J mice, the main drivers of the Steap3 signal in the previous study) that possess far greater levels of segregating genetic variants. Curse and blessing of careful and reproducible science, our results did not identify novel candidates, but rather provided an independent replication of the Steap3 finding,^77^ showing an effect on PTR and lipid peroxidation of additional strain-specific polymorphisms beyond those we had previously observed in FVB mice. Through genetic manipulation of poor and good “storer” mouse strains^27^, we show that ablation of STEAP3 or knock in of the gain of function STEAP3 are sufficient to affect lipid peroxidation and extravascular hemolysis of stored and transfused RBCs in vivo. Given the solidity and reproducibility of this finding in mice, we thus expanded on their translational relevance in humans, by showing for the first time that common polymorphisms – especially prevalent in blood donors of African descent – are associated with lower RBC iron levels, decreased lipid peroxidation and hemolysis.

Mechanistically, these data are consistent with a role for STEAP3 in promoting Fenton and Haber-Weiss chemistry in iron-loaded mature erythrocytes. Specifically, ferrous iron participates in this chemistry, becoming oxidized to its ferric state in the process with the concomitant generation of hydroxyl or hydroperoxyl radicals, which can in turn attack fatty acids (preferentially, poly and highly-unsaturated fatty acids such as octadecadienoic and eicosatetranoic acid) thus promoting lipid peroxidation. By reducing Fe^3+^ to its Fe^2+^ state, the ferrireductase STEAP3 shifts the balance of this reaction by increasing the availability of the reactant, thus favoring the generation of product radicals –consistent with ferroptosis^40^, a process also referred to as death by lipid peroxidation^41^ and that closely resembles the hallmarks of non-apoptotic RBC death or eryptosis^78^. While some of these processes have been reported to occur during RBC aging^79^, and pharmacological and genetic manipulation of ferroptosis pathways has been tested in cancer^40^, no study to date has pursued these strategies to alter RBC lifespan in vitro (blood bank storage) and in vivo. Of note, the burgeoning literature on ferroptosis has hitherto disregarded anucleate RBCs, despite two thirds of bodily iron being stored in the 25 trillion circulating erythrocytes.

Relevant to the present study, ferroptosis is promoted by TP53^47,80^ and STEAP3 in cancer^81,82^, but never hitherto described in the context of RBC aging in vivo and in vitro. Here we show that common cancer predisposing TP53 polymorphisms are not only observed in blood donors with frequencies consistent with those reported in the literature for cancer-focused studies (∼1:2,000^51^), but also associated with altered lipid peroxidation, antioxidant metabolism - especially the pentose phosphate pathway, which is consistent with the reported suppressive role of TP53 on G6PD in cancer^83^ - and hemolytic propensity of stored human RBCs. We thus show that common TP53 alleles are associated with altered post-transfusion performances of transfused units, as determined by hemoglobin increments in thousands of single unit recipients of units from donors carrying mutant TP53 alleles.

Finally, we identified novel genetic variants associated with oxylipin levels in end of storage units from over 13 thousand human blood donor volunteers. Overall, these analyses showed that genetic regulation of oxylipin levels in humans convergence on a pathway that participates in detoxification of lipid peroxidation products via acyl-carnitine (Lands cycle)^65,84^ - regulated by LPCAT3, NADPH (regulated by G6PD), and glutathione-dependent mechanisms (regulated by GSS and GPX4). The latter enzyme has been recently identified as a marker of RBC susceptibility to hemolysis^85^, especially following oxidant insults^20^. The role of GPX4 and G6PD in ferroptosis has been described in cancer, but not in the context of RBC aging in vivo and in vitro^86,87^. Elongation (ELOVL1) and desaturation (FADS1/2) of fatty acids provides the substrate for the generation of oxylipins. Only recently, fatty acid desaturases have been identified and shown to be active in mature RBCs as a function of oxidant stress and G6PD status^88^. While STEAP3 itself did not emerge from the mQTL analysis in humans, the role of STEAP3 in regulating RBC iron levels, fueling lipid peroxidation and hemolysis contributes to painting an overall coherent story between the murine and human data. In particular, we observed that the allele frequency for the TP53 SNP rs8064946 was associated with that of all the other significant SNPs from the human mQTL analysis, bridging the TP53-STEAP3 axis with the genetic traits linked to lipid peroxidation in humans. Of note, our ethnic breakdown of allele frequency for significant polymorphisms associated to oxylipins levels (STEAP3, TP53, LPCAT3, FADS1/2, EPHX2) shows an enrichment of beneficial SNPs and under-representation of deleterious SNPs associated with lipid peroxidation in donors of African descent, providing a theoretical mechanistic basis underpinning the observed, yet unexplained resistance to osmotic insults of RBCs from individuals in this ethnic group^18^.

While STEAP3 is a new potential regulator of red blood cell ferroptosis, many of the other genes we identified are already validated, key regulators of ferroptosis. The p53 tumor suppressor protein has been shown to trigger ferroptosis through downregulation of SLC7A11^89^ and through an ALOX12 pathway^90^. G6PD is a key regulator of NADPH biosynthesis, which was identified as a biomarker predicting sensitivity to ferroptosis across the NCI60 panel^91^. LPCAT3 was discovered in a genetic screen in KBM7 cells for genes whose disruption cause resistance to GPX4 inhibitors^92^. More recently, the related MBOAT genes have been discovered as key inhibitors of ferroptosis^93^. GPX4 was one of the first key regulators of ferroptosis discovered^94^. FADS1/2 and ELOVs have also been linked to ferroptosis^95,96^, along with GSH biosynthesis^97^. PLA2G6 was also identified as a key ferroptosis inhibitor^98^. These unbiased genetic results therefore strongly suggest that storage of red blood cells leads to their loss through ferroptosis, and that additional genes known to regulate ferroptosis might also control RBC stability.

### Limitations of the study

The combination of GWAS results in stored RBCs from mouse and human studies here provides complementary information, to the extent this integrated analysis allowed us to interrogate the same fundamental biology against two different collections of genetic variants from diverse murine and human populations. In this setting, blindness to signals resulting from the lack of genetic variance for a given trait in one species is overcome by genetic variance in the other species. Results from studies in mice offer a deeper look into genes and processes that are relevant in human, and vice versa, while the combination of signals from both species, while note entirely overlapping around a specific gene, ultimately does converge onto the same pathway. Specifically, altogether our data highlights the centrality of the oxidant stress-induced effect on RBC membrane lipids and enzymes’ roles in detoxification to RBC susceptibility to intra- and extra-vascular hemolysis. These results ultimately suggest a role for ferroptosis^40^ in RBC intra- and extra-vascular hemolysis, while identifying potential new regulatory players in this iron-dependent mechanism of cell death by lipid peroxidation. Understanding that the storage lesion process is a ferroptotic one has direct medical relevance. Multiple pharmacological or dietary interventions can trigger (e.g., erastin^99^), mitigate or even prevent ferroptosis in other systems, including iron chelators, inhibitors of lipid peroxidation or lipophilic antioxidant^100,101^, or dietary supplementation of omega-3 or deuterium-labeled^102^ fatty acid-enriched diets. The efficacy of these drugs in the context of blood storage remains untested and holds the potential to extend the shelf-life, quality and post-transfusion performances of packed RBC products, as well as paving the way for new treatment strategies for hemolytic disorders. Finally, with the advent of the era of Precision Transfusion Medicine, the findings reported herein can inform on genetic screening for alleles (e.g., STEAP3, TP53, LPCAT3) that are associated with poorer/improved storage quality, with the goal to inform shortening/extension of the shelf-life of units from donors carrying these polymorphisms, to guide priority in blood unit issuing based on genetically-informed prediction of storability rather than a first-in/first-out strategy in the blood bank.

## STAR METHODS

### RESOURCE AVAILABILITY

#### Lead contact

Further information and requests for resources and reagents should be directed to and will be fulfilled by the Lead Contact, Angelo D’Alessandro

#### Materials availability

No new materials were generated as part of this study. Information on the human and murine samples tested in this study is detailed below.

#### Data and code availability

The mouse data (metabolites, lipids, oxylipins, PTR, genotypes) and R code for processing data and producing results have been made publicly available at figshare (https://doi.org/10.6084/m9.figshare.24456619.v1). Processed data and results can also be downloaded from the QTLViewer (https://churchilllab.jax.org/qtlviewer/Zimring/RBC). All the raw data and elaborations are provided in **Supplementary Table 1**.

### EXPERIMENTAL MODEL AND SUBJECT DETAILS

#### J:DO mouse studies: RBC storage and post-transfusion recovery

Mouse storage were performed as previously described ^31^. A total of 350 J:DO mice were derived from extensive outbreeding of eight inbred founder strains that represent genetically distinct lineages of the house mouse: A/J, C57BL/6J, 129S1/SvlmJ, NOD/ShiLtJ, NZO/HILtJ, CAST/EiJ, PWK/PhJ, WSB/EiJ (**Figure 1.A**). All animal procedures were approved by the University of Virginia IACUC (protocol no. 4269). All animals were genotyped at 143,259 SNPs using the GigaMUGA array ^32^.

#### Post-transfusion recovery (PTR) studies in mice

Mouse post-transfusion recovery (PTR) studies were performed as previously described ^103^. Storage of diversity outbred mouse RBCs (n=349, owing to technical issues with one tube for one of original 350 mice) for 7 days was followed by transfusion into Ubi-GFP mice, which were used as recipients to allow visualization of the test cells in the non-fluorescent gate. To control for differences in transfusion and phlebotomy, biotinylated RBCs were used as a tracer RBC population (never stored). These RBCs were added to stored RBCs immediately prior to transfusion. PTR was calculated by dividing the post-transfusion ratio (Test/Tracer) by the pre-transfusion ratio (Test/Tracer), with the maximum value set as 1 (or 100% PTR).

#### QTL analysis in JAX DO mice

The QTL workflow (for PTR, mQTL and lQTL) in JAX DO mice followed previously defined conventions ^104^. Briefly, metabolite values of zero were converted to a missing value and only metabolites or lipids with 100 non-missing observations or more were included for further analyses. Each metabolite was transformed to normal quantiles for mQTL and lQTL analysis to reduce the influence of outlying values. The initial mQTL and lQTL mapping was based on founder strain haplotypes imputed at 137,192 loci, allowing for the additive genetic effects at a locus to be more richly characterized as eight founder allele effects. Sex (148 females and 202 males) and sub-cohorts (5 groups ranging from 27 to 94 mice) were adjusted for as covariates. A lenient significance threshold of LOD score > 7 was used for calling mQTL/lQTL to allow for the detection of mQTL/lQTL hotspots. For reference, a LOD score > 8 approximately represents a genome-wide adjusted p-value < 0.05 ^105^. Fine mapping at detected mQTL/lQTL was performed by imputing variants based on the genotypes of the founder strains (GRCm39). We used the same mQTL/lQTL workflow for fresh and stored RBCs. All analyses were performed using the R package qtl2 ^104^.

#### Donor recruitment in the REDS RBC Omics study

##### Index donors

A total of 13,758 donors were enrolled in the Recipient Epidemiology and Donor evaluation Study (REDS) RBC Omics at four different blood centers across the United States (https://biolincc.nhlbi.nih.gov/studies/reds_iii/). Of these, 97% (13,403) provided informed consent and 13,091 were available for metabolomics analyses in this study – henceforth referred to as “index donors”. A subset of these donors were evaluable for hemolysis parameters, including spontaneous (n=12,753) and stress (oxidative and osmotic) hemolysis analysis (n=10,476 and 12,799, respectively) in ∼42-day stored leukocyte-filtered packed RBCs derived from whole blood donations from this cohort ^14^. Methods for the determination of FDA-standard spontaneous (storage) hemolysis test, osmotic hemolysis (pink test) and oxidative hemolysis upon challenge with AAPH have been extensively described elsewhere ^18^.

##### Recalled donors

A total of 643 donors scoring in the 5^th^ and 95^th^ percentile for hemolysis parameters at the index phase of the study were invited to donate a second unit of pRBCs, a cohort henceforth referred to as “recalled donors”. These units were assayed at storage days 10, 23 and 42 for hemolytic parameters and mass spectrometry-based high-throughput metabolomics ^24^, proteomics ^106^, lipidomics ^35^ and ICP-MS analyses ^107^. Under the aegis of the REDS-IV-P project ^108^, a total of 1,929 samples (n=643, storage day 10, 23 and 42) were processed with this multi-omics workflow.

### METHOD DETAILS

#### High-throughput metabolomics

Metabolomics extraction and analyses in 96 well-plate format were performed as described ^109,110^. RBC samples were transferred on ice on 96 well plate and frozen at -80 °C at Vitalant San Francisco prior to shipment in dry ice to the University of Colorado Anschutz Medical Campus. Plates were thawed on ice then a 10 uL aliquot was transferred with a multi-channel pipettor to 96-well extraction plates. A volume of 90 uL of ice cold 5:3:2 MeOH:MeCN:water (*v/v/v*) was added to each well, with an electronically-assisted cycle of sample mixing repeated three times. Extracts were transferred to 0.2 µm filter plates (Biotage) and insoluble material was removed under positive pressure using nitrogen applied via a 96-well plate manifold. Filtered extracts were transferred to an ultra-high-pressure liquid chromatography (UHPLC-MS — Vanquish) equipped with a plate charger. A blank containing a mix of standards detailed before ^111^ and a quality control sample (the same across all plates) were injected 2 or 5 times each per plate, respectively, and used to monitor instrument performance throughout the analysis. Metabolites were resolved on a Phenomenex Kinetex C18 column (2.1 x 30 mm, 1.7 um) at 45 °C using a 1-minute ballistic gradient method in positive and negative ion modes (separate runs) over the scan range 65- 975 m/z exactly as previously described.^109^ The UHPLC was coupled online to a Q Exactive mass spectrometer (Thermo Fisher). The Q Exactive MS was operated in negative ion mode, scanning in Full MS mode (2 μscans) from 90 to 900 m/z at 70,000 resolution, with 4 kV spray voltage, 45 sheath gas, 15 auxiliary gas. Following data acquisition, .raw files were converted to .mzXML using RawConverter then metabolites assigned and peaks integrated using ElMaven (Elucidata) in conjunction with an in-house standard library ^112^.

#### High-throughput lipidomics

For lipidomics analyses, extraction procedures were identical to those described for metabolomics, though pure methanol was used. Lipid extracts were analyzed (10 µL per injection) on a Thermo Vanquish UHPLC/Q Exactive MS system using a previously described^35^ 5 min lipidomics gradient and a Kinetex C18 column (30 x 2.1 mm, 1.7 µm, Phenomenex) held at 50 °C. Mobile phase A: 25:75 MeCN:water with 5 mM ammonium acetate; Mobile phase B: 90:10 isopropanol:MeCN with 5 mM ammonium acetate. The gradient and flow rate were as follows: 0.3 mL/min of 10% B at 0 min, 0.3 mL/min of 95% B at 3 min, 0.3 mL/min of 95% B at 4.2 min, 0.45 mL/min 10% B at 4.3 min, 0.4 mL/min of 10% B at 4.9 min, and 0.3 mL/min of 10% B at 5 min. Samples were run in positive and negative ion modes (both ESI, separate runs) at 125 to 1500 m/z and 70,000 resolution, 4 kV spray voltage, 45 sheath gas, 25 auxiliary gas. The MS was run in data-dependent acquisition mode (ddMS^2^) with top10 fragmentation. Raw MS data files were searched using LipidSearch v 5.0 (Thermo).

#### REDS RBC Omics mQTL analyses of oxylipins

The workflow for the mQTL analysis of oxylipin metabolites is consistent with previously described methods from our pilot mQTL study on 250 recalled donors ^22^. Details of the genotyping and imputation of the RBC Omics study participants have been previously described by Page, et al. ^20^ Briefly, genotyping was performed using a Transfusion Medicine microarray ^43^ consisting of 879,000 single nucleotide polymorphisms (SNPs); the data are available in dbGAP accession number phs001955.v1.p1. Imputation was performed using 811,782 SNPs that passed quality control. After phasing using Shape-IT ^113^, imputation was performed using Impute2 ^114^ with the 1000 Genomes Project phase 3 ^114^ all-ancestry reference haplotypes. We used the R package SNPRelate ^115^ to calculate principal components (PCs) of ancestry. We performed association analyses for kynurenine using an additive SNP model in the R package ProbABEL ^116^ and 13,091 study participants who had both metabolomics data and imputation data on serial samples from stored RBC components that passed respective quality control procedures. We adjusted for sex, age (continuous), frequency of blood donation in the last two years (continuous), blood donor center, and ten ancestry PCs. Statistical significance was determined using a p-value threshold of 5x10^-8^. We only considered variants with a minimum minor allele frequency of 1% and a minimum imputation quality score of 0.80. The OASIS: Omics Analysis, Search & Information a TOPMED funded resources ^117^, was used to annotate the top SNPs. OASIS annotation includes information on position, chromosome, allele frequencies, closest gene, type of variant, position relative to closest gene model, if predicted to functionally consequential, tissues specific gene expression, and other information.

#### Determination of hemoglobin and bilirubin increment via the vein-to-vein database

Association of metabolite levels with hemoglobin increments was performed by interrogating the vein-to-vein database, as described in Roubinian et al. ^17,21^

#### Vein-to-vein database: General Study Design

We conducted a retrospective cohort study using electronic health records from the National Heart Lung and Blood Institute (NHLBI) Recipient Epidemiology and Donor Evaluation Study-III (REDS-III) program available as public use data through BioLINCC ^118,119^. The database includes blood donor, component manufacturing, and patient data collected at 12 academic and community hospitals from four geographically diverse regions in the United States (Connecticut, Pennsylvania, Wisconsin, and California) for the 4-year period from January 1, 2013 to December 31, 2016. Genotype and metabolomic data from the subset of blood donors who participated in the REDS-III RBC-Omics study ^120^ was linked to the dataset using unique donor identifiers.

#### Study Population and Definitions

Available donor genetic polymorphism and metabolomic data was linked to issued RBC units using random donor identification numbers. Among transfusion recipients, we included all adult patients who received a single RBC unit during one or more transfusion episodes between January 1, 2013 and December 30, 2016. Recipient details included pRBC storage age, and blood product issue date and time. We collected hemoglobin levels measured by the corresponding clinical laboratory prior to and following each RBC transfusion event (0h and 24h after transfusion).

#### Transfusion Exposures and Outcome Measures

All single RBC unit transfusion episodes linked to genetic polymorphism and metabolomic data were included in this analysis. A RBC unit transfusion episode was defined as any single RBC transfusion from a single donor with both informative pre- and post-transfusion laboratory measures and without any other RBC units transfused in the following 24-hour time period. The outcome measures of interest were change in hemoglobin (ΔHb; g/dL) following a single RBC unit transfusion episode. These outcomes were defined as the difference between the post-transfusion and pre-transfusion levels. Pre-transfusion thresholds for these measures were chosen to limit patient confounding (e.g., underlying hepatic disease). For pre-transfusion hemoglobin, the value used was the most proximal hemoglobin measurements prior to RBC transfusion, but at most 24 hours prior to transfusion. Furthermore, we excluded transfusion episodes where the pre-transfusion hemoglobin was greater than 9.5 g/dL, and the hemoglobin increment may be confounded by hemorrhage events. For post-transfusion hemoglobin, the laboratory measure nearest to 24-hours post-transfusion, but between 12- and 36- hours following transfusion was used.

### QUANTIFICATION AND STATISTICAL ANALYSIS

#### Statistical Methods

We assessed the univariable association of all *a priori* selected donor, manufacturing, and recipient covariates with the outcomes using linear regression. Multivariable linear regression assessed associations between alleles of donor single nucleotide polymorphisms and changes in hemoglobin levels post-transfusion hemoglobin. Two-sided p-values less than 0.05 were considered to be statistically significant. Analyses were performed using Stata Version 14.1, StataCorp, College Station, TX.

#### Data analysis

Statistical analyses – including hierarchical clustering analysis (HCA), linear discriminant analysis (LDA), uniform Manifold Approximation and Projection (uMAP), correlation analyses and Lasso regression were performed using both MetaboAnalyst 5.0 ^121^ and in-house developed code in R (4.2.3 2023-03-15).

## Acknowledgements

AD and JCZ were supported by funds by the National Heart, Lung, and Blood Institute (NHLBI) (R21HL150032, R01HL146442, R01HL149714, R01HL148151). The REDS RBC Omics and REDS-IV-P CTLS programs are sponsored by the NHLBI contract 75N2019D00033, and from the NHLBI Recipient Epidemiology and Donor Evaluation Study-III (REDS-III) RBC Omics project, which was supported by NHLBI contracts HHSN2682011-00001I, -00002I, -00003I, -00004I, -00005I, -00006I, -00007I, -00008I, and -00009I. B.R.S. was supported by National Cancer Institute (NCI) grant R35CA209896. G.R.K and G.A.C were supported by grants from the National Institute of General Medical Sciences (NIGMS), F32GM124599 and R01GM067945, respectively. NR received funding from NHLBI (R01HL126130). The content is solely the responsibility of the authors and does not necessarily represent the official views of the National Institutes of Health. We thank Corinne M. Keele for her illustrations of the J:DO founder strains (**Figure 1.A**).

## Authorship

Contribution: Animal studies: AH, JCZ. Metabolomics analyses: TN, DS, AD. Proteomics: MD, KCH. Biostatistics and Bioinformatics: GRK, GAC, XD, AD. REDS RBC Omics: MS, SK, SLS, PJN, MPB. Vein-to-vein database: NR. mQTL analyses: GRK, GAC (mouse) EA, GPP (human). Figure preparation: GAC, EA, AD. Writing: AD.

