## Supplementary Figures for "Ferroptosis regulates hemolysis in stored murine and human red blood cells"

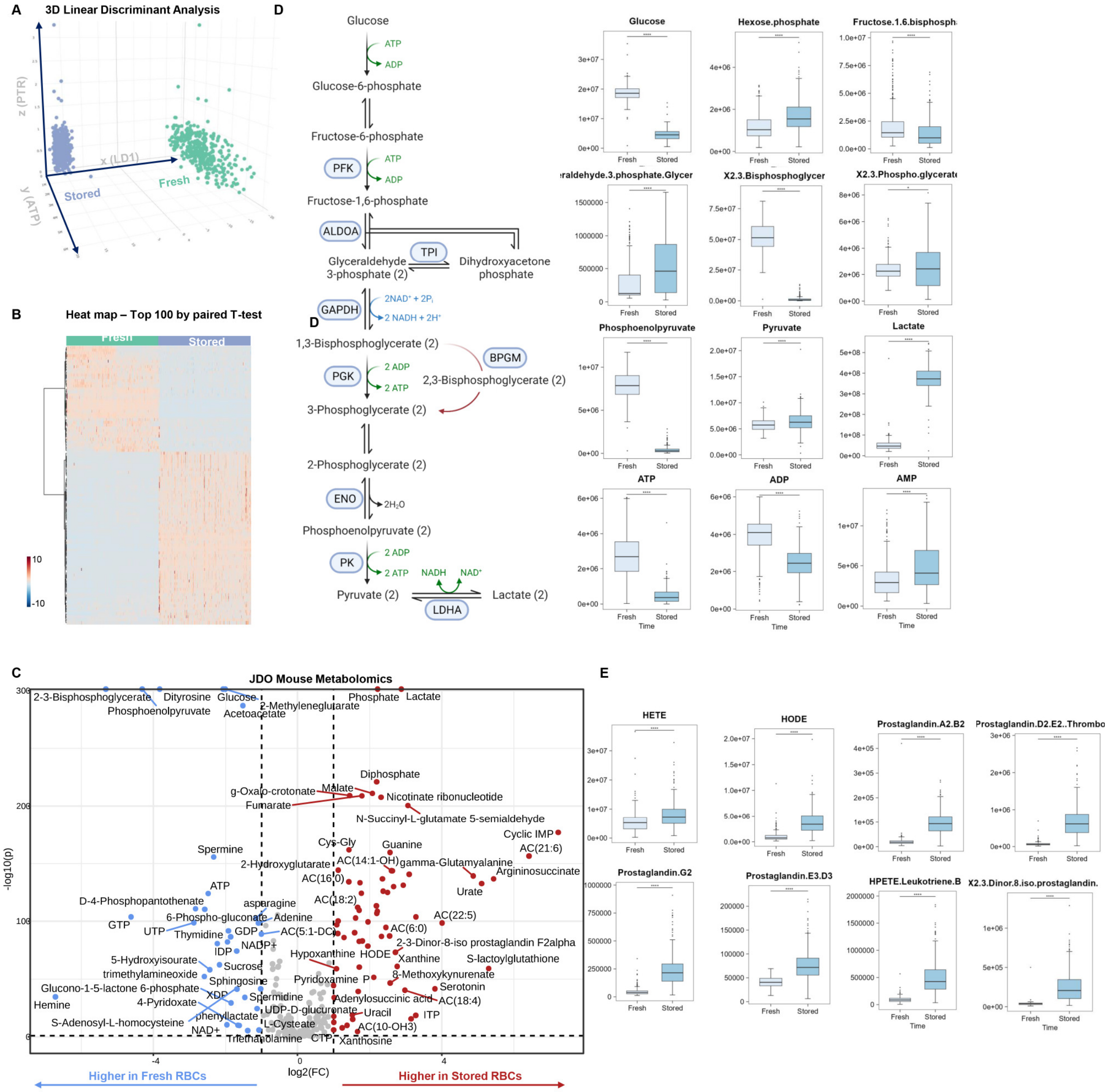


**Supplementary Figure 1 – Metabolic markers of the storage lesion in the JAX Diversity Outbred (JDO) mice.** Eight genetically diverse founder strains underwent extensive cross-breeding to obtain 350 genetically diverse mice. RBCs from these mice were stored under conditions mimicking human RBC storage in the blood bank for 7 days (equivalent to day 42 in humans). Metabolomics analyses in these 350 mice (x2 time points, day 0 fresh and day 7 stored samples) showed a significant effect of storage duration, as gleaned by linear-discriminant analysis (**A**) and hierarchical clustering analysis of the top 50 metabolites by repeated measures ANOVA (**B**) and volcano plot analyses (**C**). Markers included a significant alteration of glycolysis (**D**), the only energy generating pathway in mature RBCs, and lipid peroxidation markers (**E**).

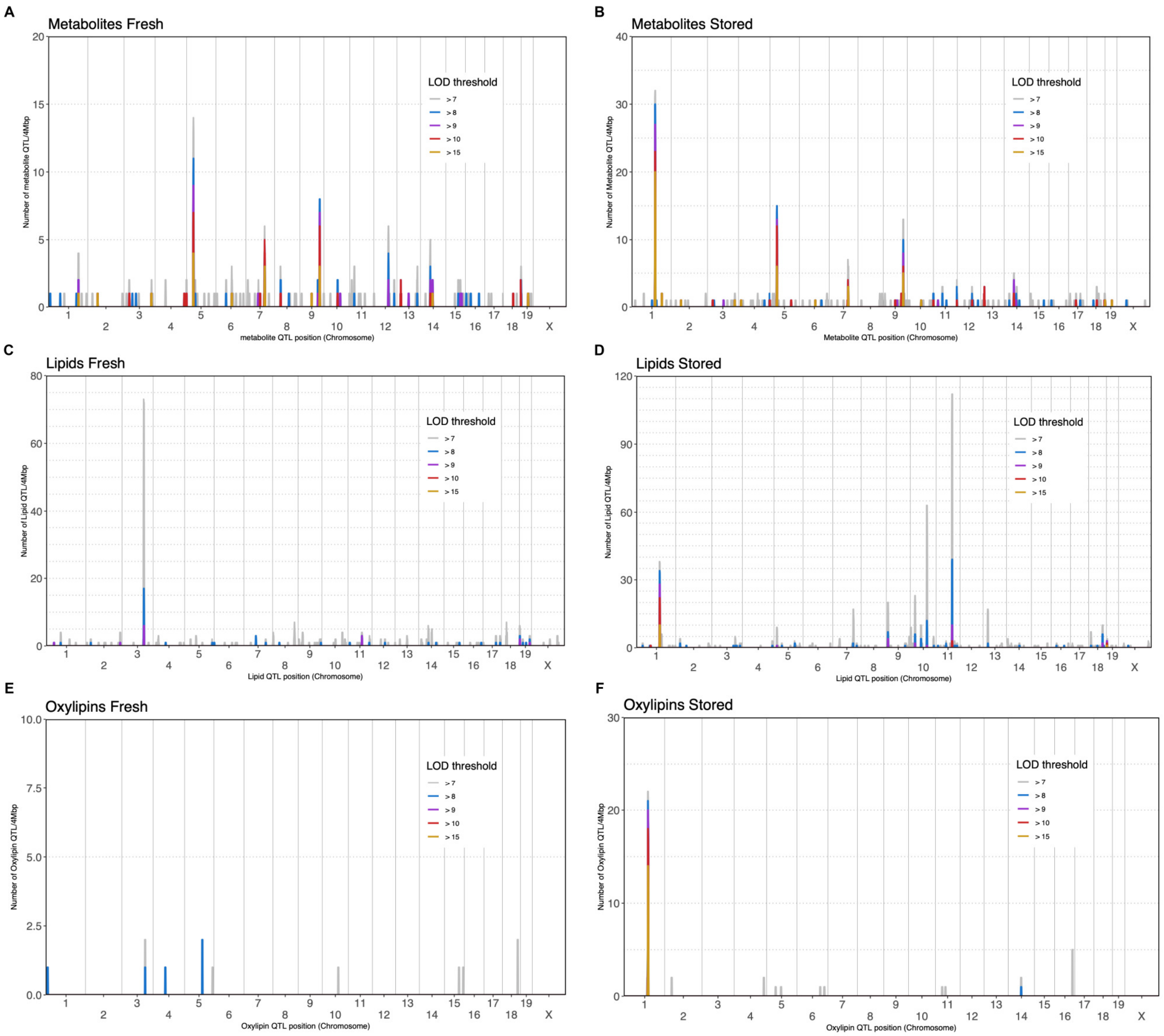


**Supplementary Figure 2 – QTL density plots for metabolites (A-B), lipids (C-D) and oxylipins (E-F).** Results from fresh data are in the left column and stored data in the right column.

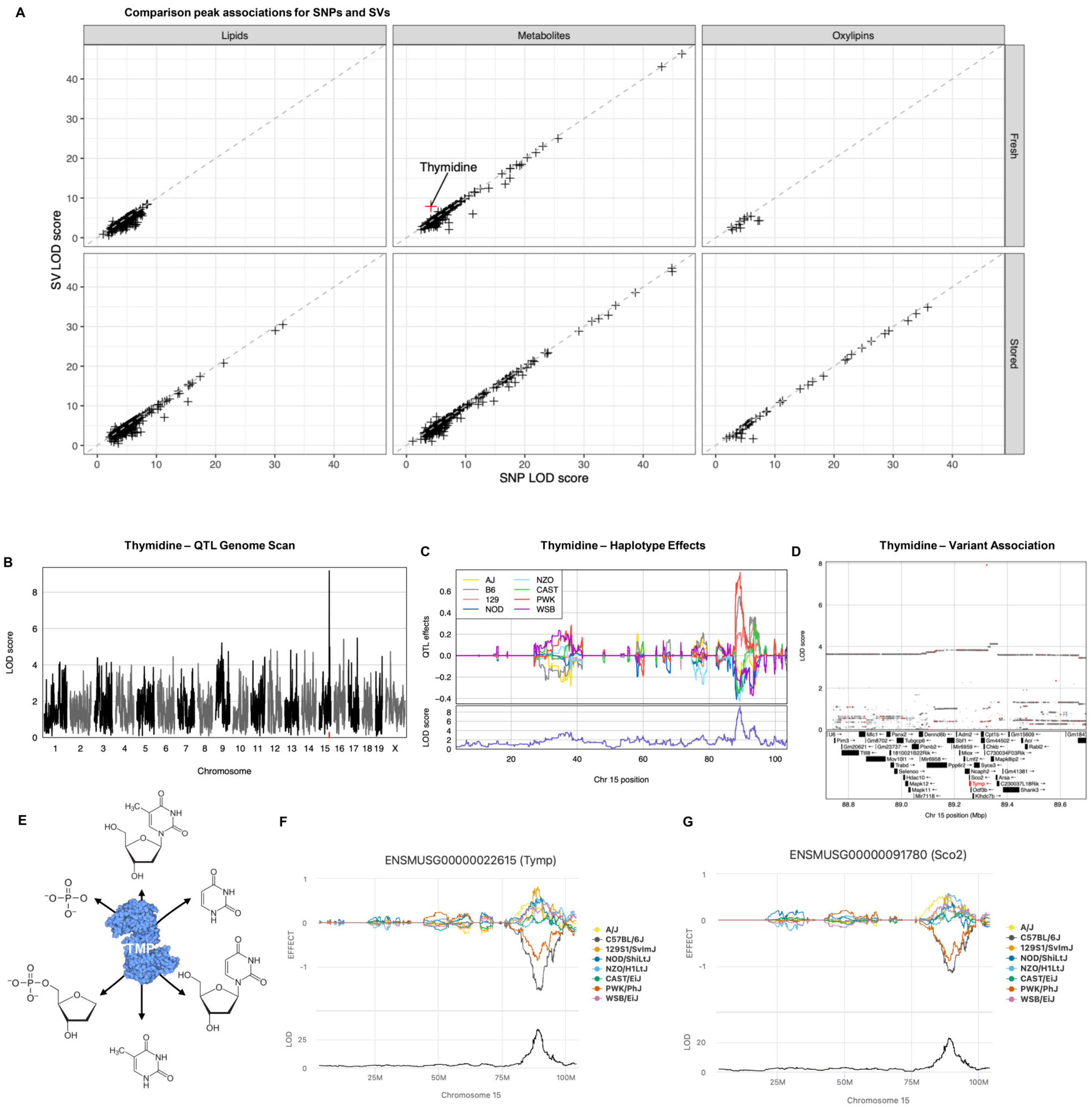


**Supplementary Figure 3 – Leveraging systems genetics to fine-map QTL for thymidine levels.** Comparison of peak SNP and structural variant (SV) associations at detected QTL for lipids, metabolites and oxylipins from fresh and stored data (**A**). The QTL for thymidine is on chromosome 15 (**B**) and is characterized by high B6 and PWK haplotype effects (**C**). The peak SV is located nearby upstream of the gene thymidine phosphorylase (TYMP – **D**). Overview of the metabolic reactions catalyzed by Tymp (**E**). In liver tissue from another DO cohort, expression QTL (eQTL) were detected for Tymp (**F**) as well as Sco2 (**G**), with inverted haplotype effects (low B6 and PWK). These effects are consistent with the SV disrupting expression levels of Tymp and Sco2, thus resulting in higher levels of thymidine due to reduction in its metabolism.

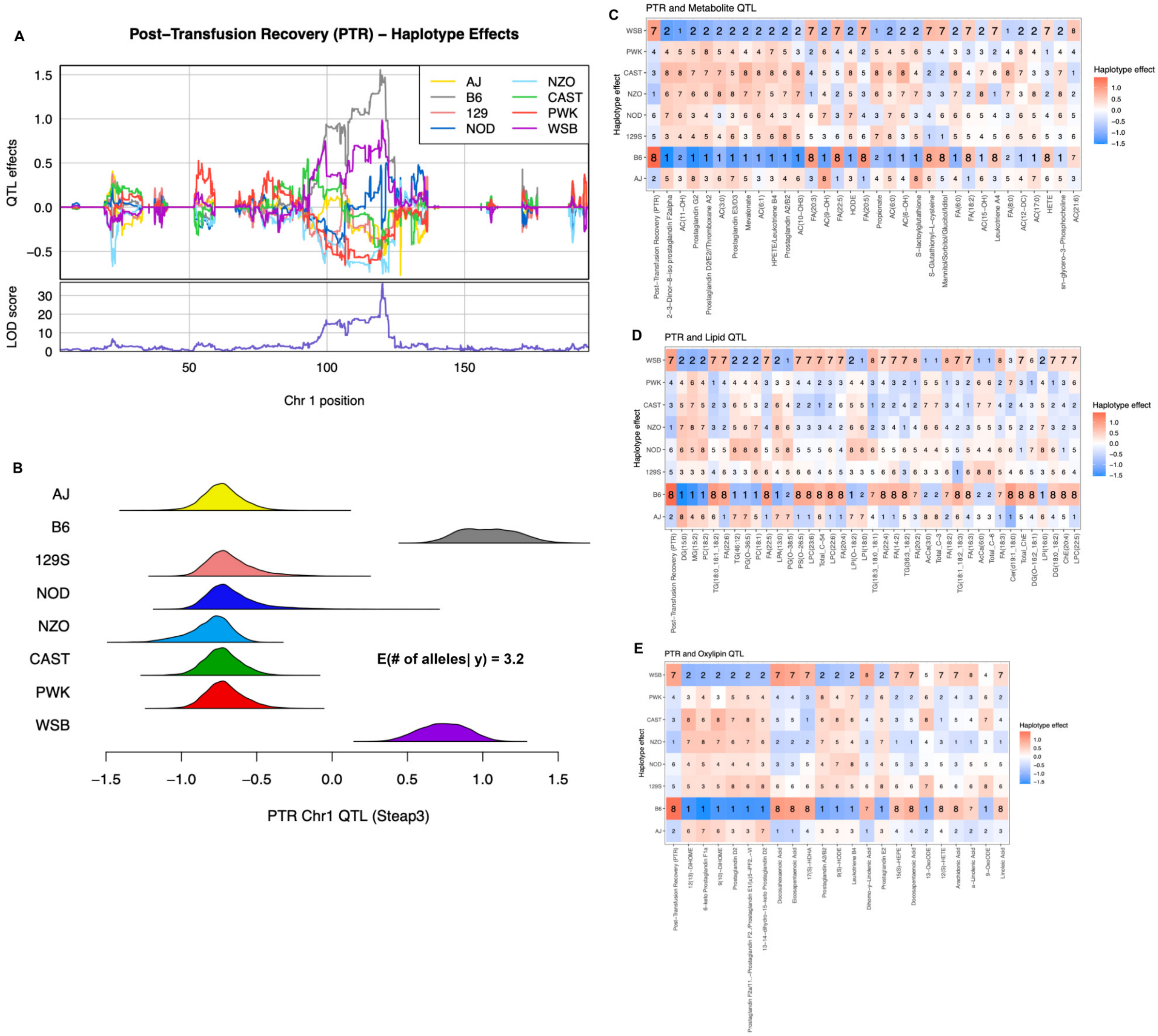


**Supplementary Figure 4 – Support for multiple functional alleles at Steap3 QTL.** Genome scan and haplotype effects at the Steap3 locus for post-transfusion recovery (PTR – **A**). Posterior effect estimates from TIMBR, a Bayesian method for modeling allelic series, estimates the posterior number of functional alleles as 3.2, with the highest posterior allelic series of c([B6], [WSB], [AJ, 129S, NOD, NZO, CAST, PWK]), suggesting that the B6 and WSB effects represent functionally distinct alleles. Ranking the haplotype effects at QTL for PTR, metabolites (**C**), lipids (**D**) and oxylipins (**E**). Numbers in cells represent the haplotype effect rank for a given QTL mapping to the Steap3 locus. Rankings of 1 and 8 for B6 and 2 and 7 for WSB are in bold to highlight the consistency of the haplotype effects.

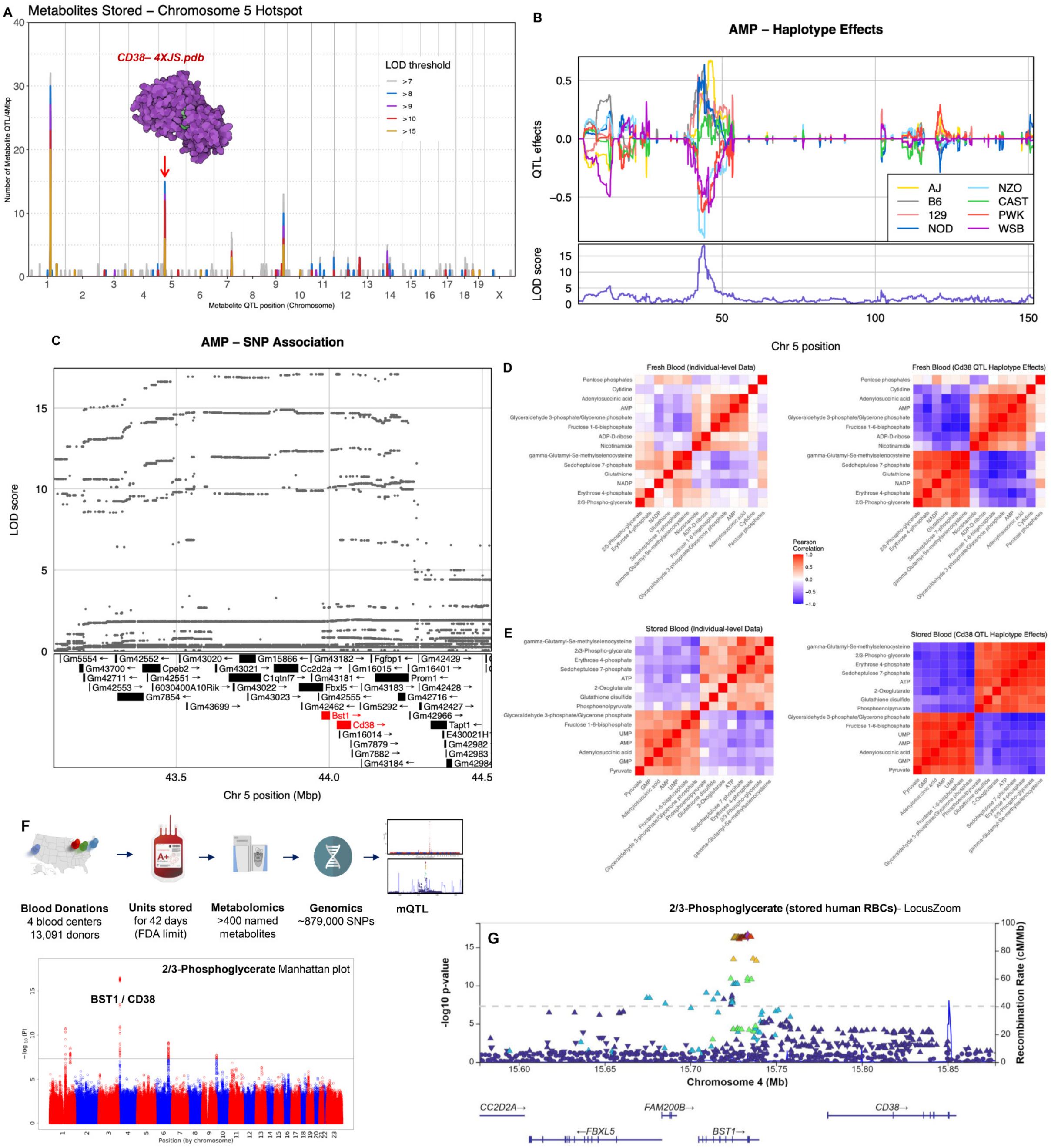


**Supplementary Figure 5 – Chromosome 5 QTL hotspot at the CD38/BST1 locus is associated with glycolytic metabolites in mice and humans.** QTL density plot for metabolites in stored samples (**A**). Chromosome 5 hotspot is also observed for metabolites in fresh samples (**Figure S1A**). Genome scan and haplotype effects at the Cd38/Bst1 locus for adenosine monophosphate (AMP – **B**). SNP associations at AMP QTL and genes in the region (**C**). The correlation structure among metabolites reflects the Cd38/Bst1 QTL, for both fresh samples (**D**) and stored samples (**E**), at both the individual-level data (left) and the QTL haplotype effects (right). In **F**, mQTL analysis in REDS RBC Omics identified CD38/BST1 as a hotspot for multiple glycolytic metabolites, including 2/3-phosphoglycerate (both isomers combined for the Manhattan plot and locus zoom analyses in **F-G**).

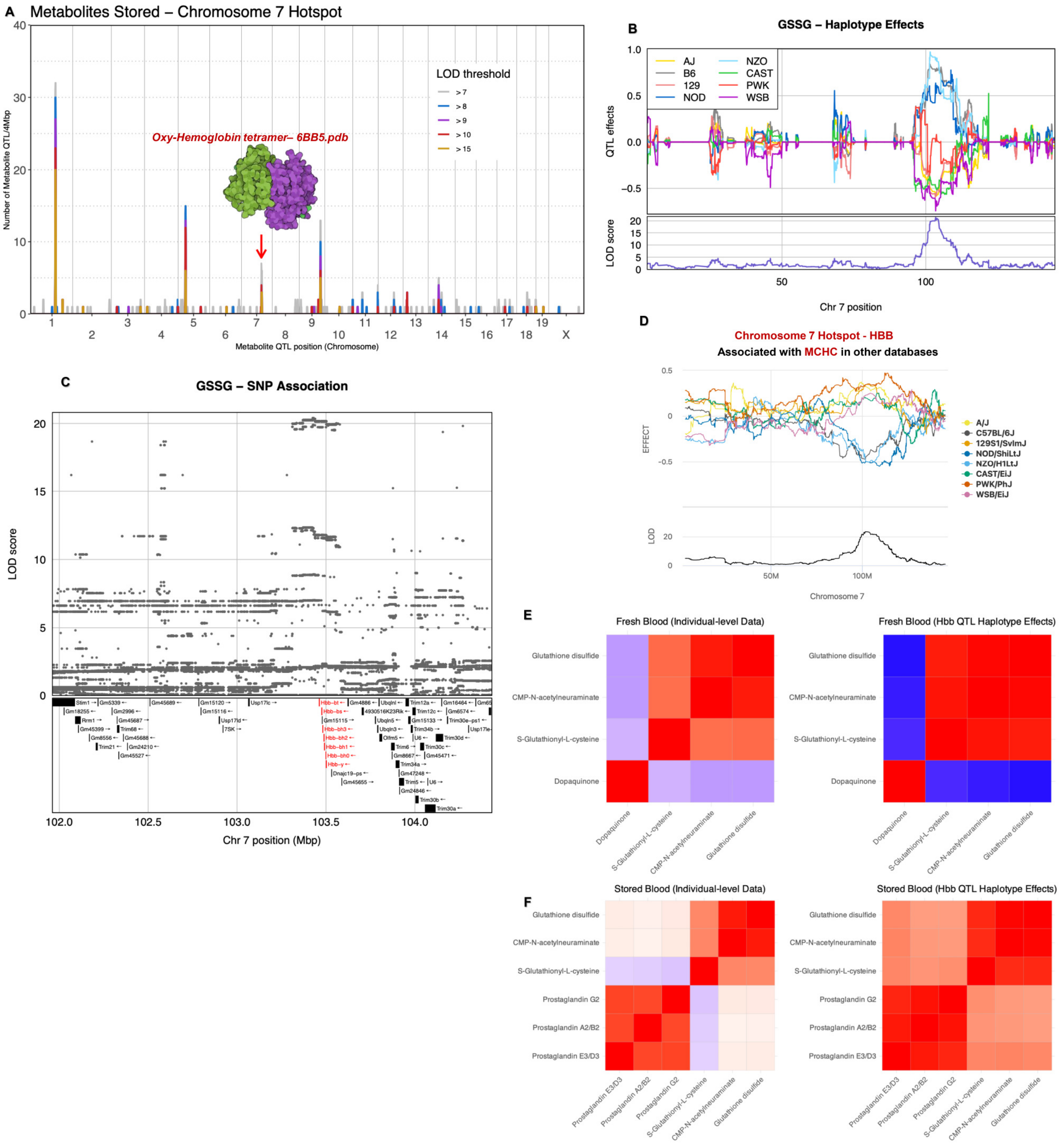


**Supplementary Figure 6 – Chromosome 7 QTL hotspot at the hemoglobin beta (Hbb) locus is associated with antioxidant metabolites and glutathione homeostasis.** QTL density plot for metabolites in stored samples (**A**). Chromosome 7 hotspot is also observed for metabolites in fresh samples (**Figure S1A**). Genome scan and haplotype effects at the Hbb locus for oxidized glutathione disulfide (GSSG – **B**). SNP associations at GSSG QTL and genes in the region (**C**). Olfactory receptor (Olfr) genes were filtered out for clarity. In complete blood count data from another DO cohort, a QTL was mapped for mean cell hemoglobin concentration (MCHC) at the Hbb locus with inverted haplotype effects in comparison to the GSSG QTL (**D**). The correlation structure among metabolites reflects the Hbb QTL, for both fresh samples (**E**) and stored samples (**F**), at both the individual-level data (left) and the QTL haplotype effects (right).

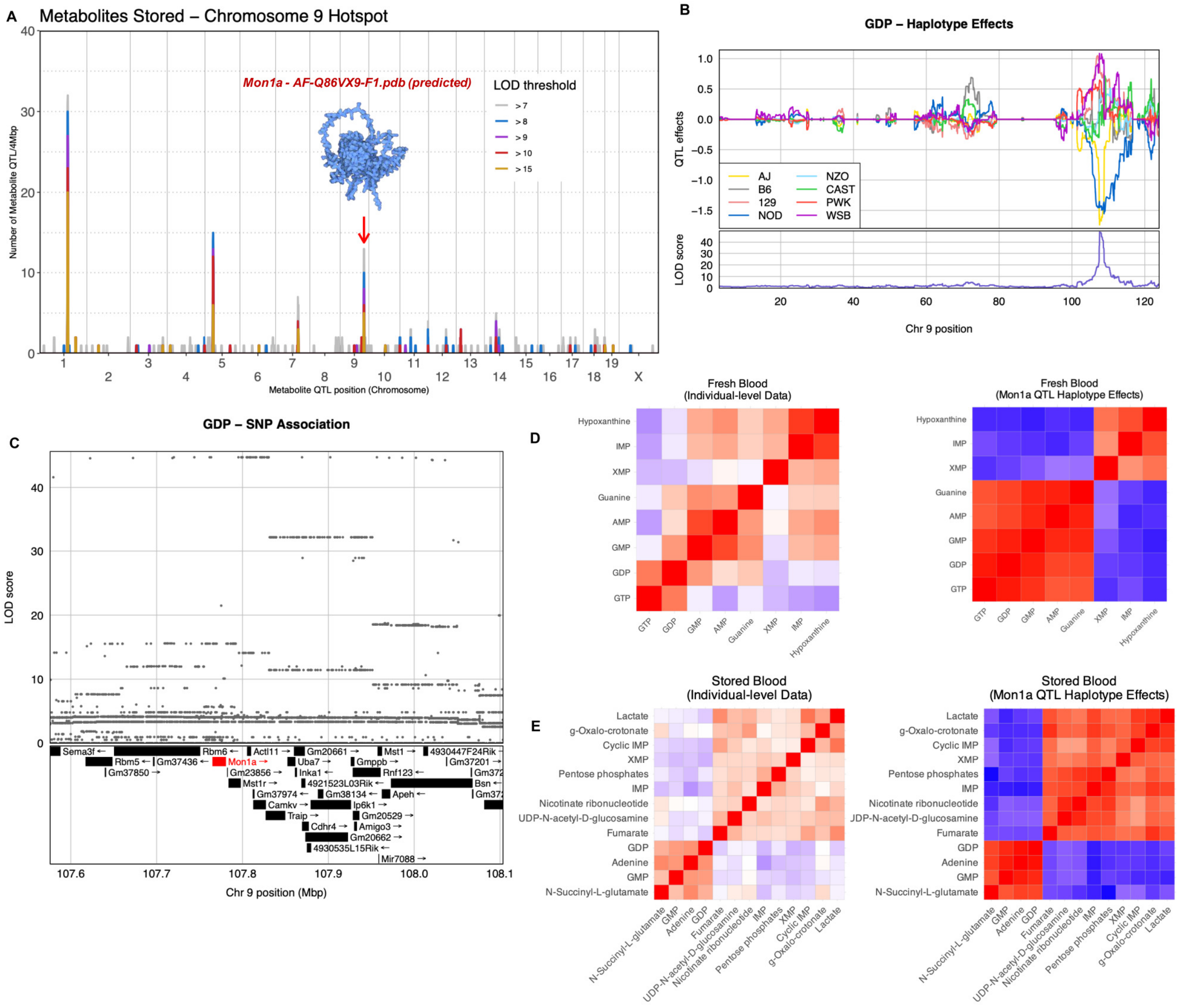


**Supplementary Figure 7 – Chromosome 9 QTL hotspot at the Mon1a locus is associated with purine monophosphate metabolites in mice.** QTL density plot for metabolites in stored samples (**A**). Chromosome 9 hotspot is also observed for metabolites in fresh samples (**Figure S1A**). Genome scan and haplotype effects at the Mon1a locus for guanosine diphosphate (GDP – **B**). SNP associations at GDP QTL and genes in the region (**C**). The correlation structure among metabolites reflects the Mon1a QTL, for both fresh samples (**D**) and stored samples (**E**), at both the individual-level data (left) and the QTL haplotype effects (right).

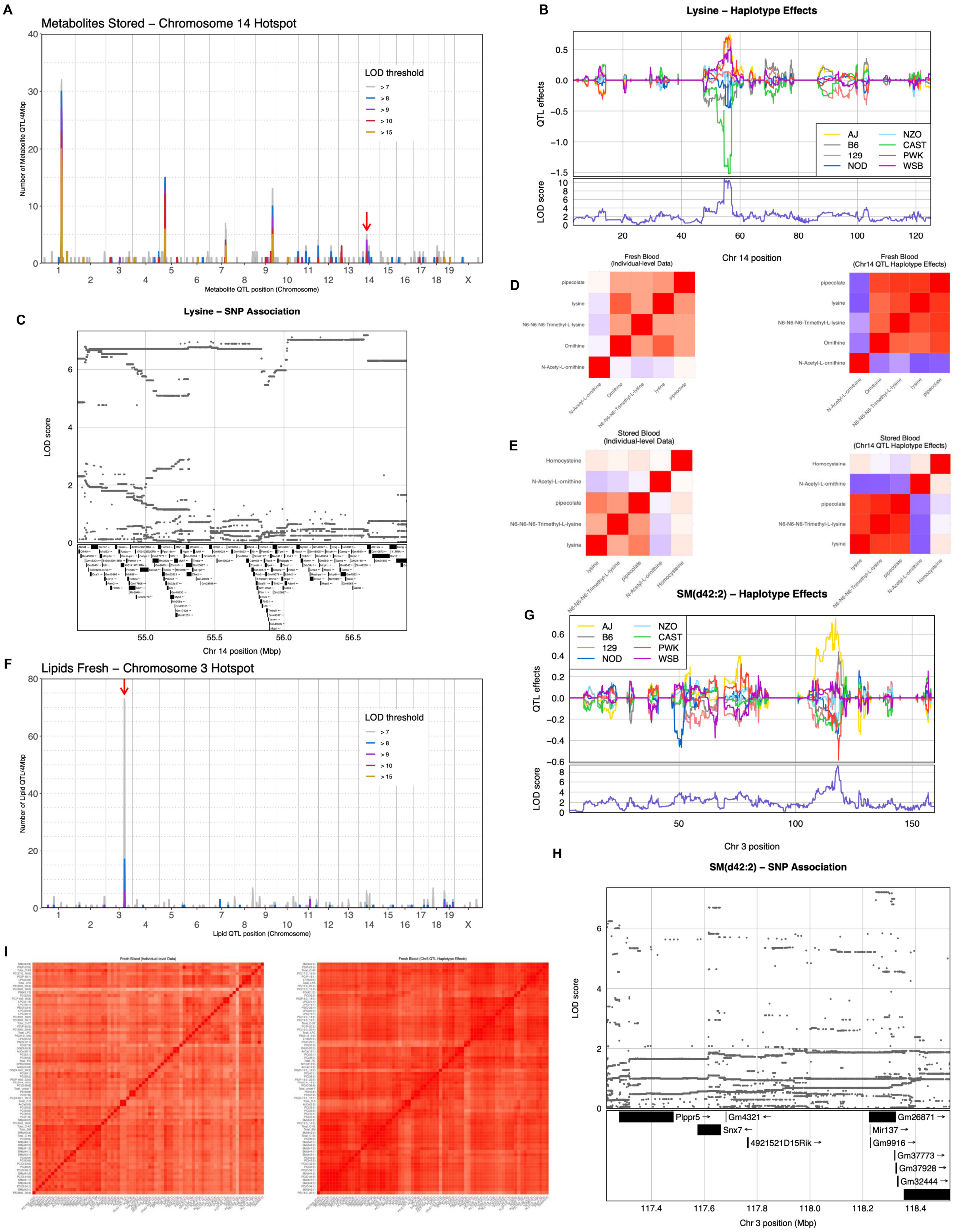


**Supplementary Figure 8 – Chromosome 14 QTL hotspot is associated with lysine metabolism and Chromosome 3 QTL hotspot is associated with lipid metabolism in mice.** QTL density plot for metabolites in stored samples (**A**). Chromosome 14 hotspot is also observed for metabolites in fresh samples (**Figure S1A**). Genome scan and haplotype effects at the hotspot locus for lysine (**B**). SNP associations at lysine QTL and genes in the region (**C**). The correlation structure among metabolites reflects the QTL hotspot, for both fresh samples (**D**) and stored samples (**E**), at both the individual-level data (left) and the QTL haplotype effects (right). QTL density plot for lipids in fresh samples (**F**). Genome scan and haplotype effects at the hotspot locus for sphingomyelin d42:2 [SM(d42:2)] (**G**). SNP associations at SM(d42:2) QTL and genes in the region (**H**). The correlation structure among lipids reflects the QTL hotspot in fresh samples (**I**), at both the individual-level data (left) and the QTL haplotype effects (right).

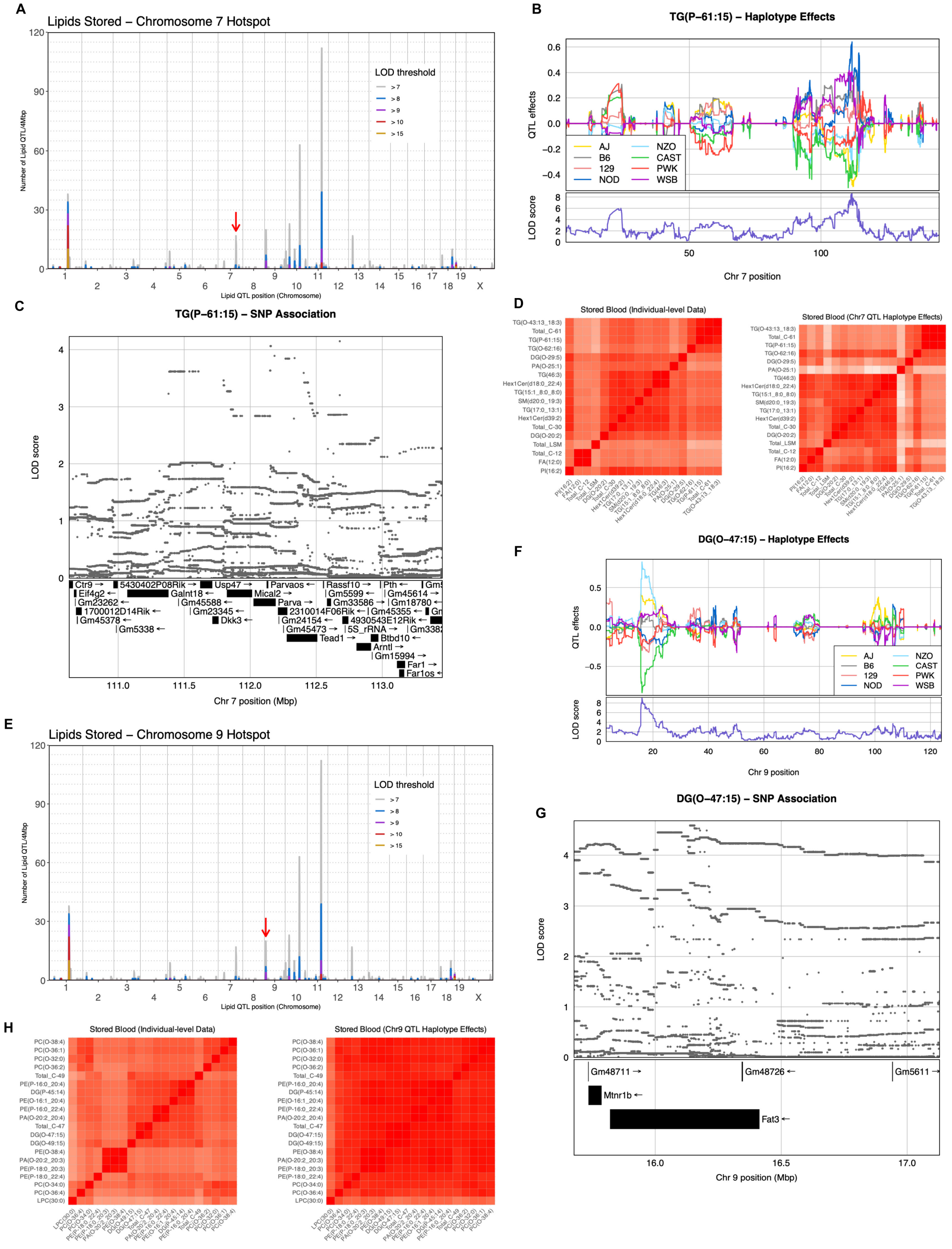


**Supplementary Figure 9 – Chromosomes 7 and 9 QTL hotspots are associated with lipid metabolism in mice.** QTL density plot for lipids in stored samples (**A, E**). Genome scan and haplotype effects at the hotspot locus for triacylglycerol P-61:15 [TG(P-61:15)] (**B**). SNP associations at TG(P-61:15) QTL and genes in the region (**C**). The correlation structure among lipids reflects the QTL hotspot in stored samples (**D**), at both the individual-level data (left) and the QTL haplotype effects (right). Genome scan and haplotype effects at the hotspot locus for diacylglycerol O-47:15 [DG(O-47:15)] (**F**). SNP associations at DG(O-47:15) QTL and genes in the region (**G**). The correlation structure among lipids reflects the QTL hotspot in stored samples (**H**), at both the individual-level data (left) and the QTL haplotype effects (right).

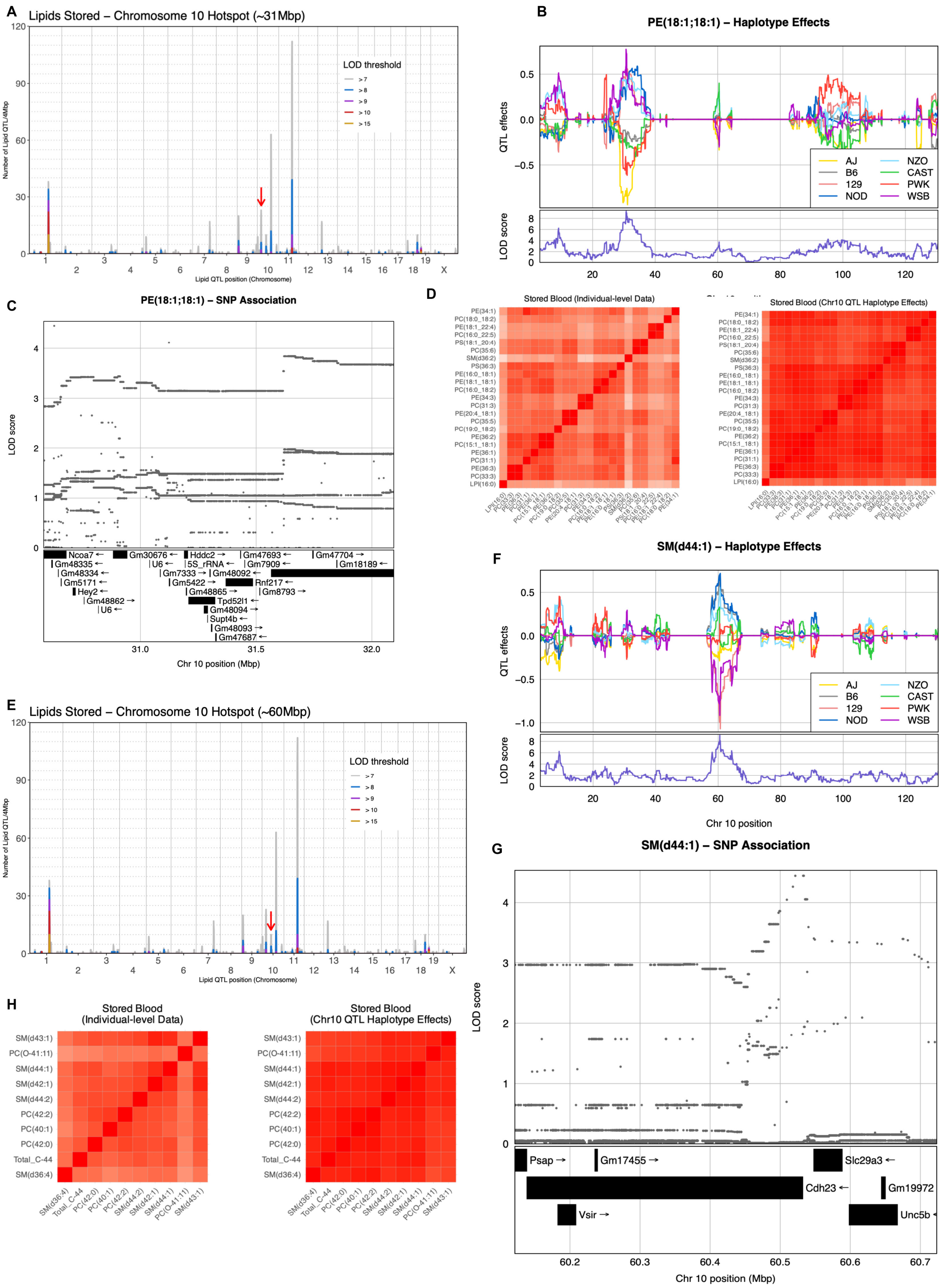


**Supplementary Figure 10 – Chromosome 10 QTL hotspots are associated with lipid metabolism in mice.** QTL density plot for lipids in stored samples (**A, E**). Genome scan and haplotype effects at the hotspot locus for phosphatidylethanolamine 18:1; 18:1 [PE(18:1;18:1)] (**B**). SNP associations at PE(18:1;18:1) QTL and genes in the region (**C**). The correlation structure among lipids reflects the QTL hotspot in stored samples (**D**), at both the individual-level data (left) and the QTL haplotype effects (right). Genome scan and haplotype effects at the hotspot locus for sphingomyelin d44:1 [SM(d44:1)] (**F**). SNP associations at SM(d44:1) QTL and genes in the region (**G**). The correlation structure among lipids reflects the QTL hotspot in stored samples (**H**), at both the individual-level data (left) and the QTL haplotype effects (right).

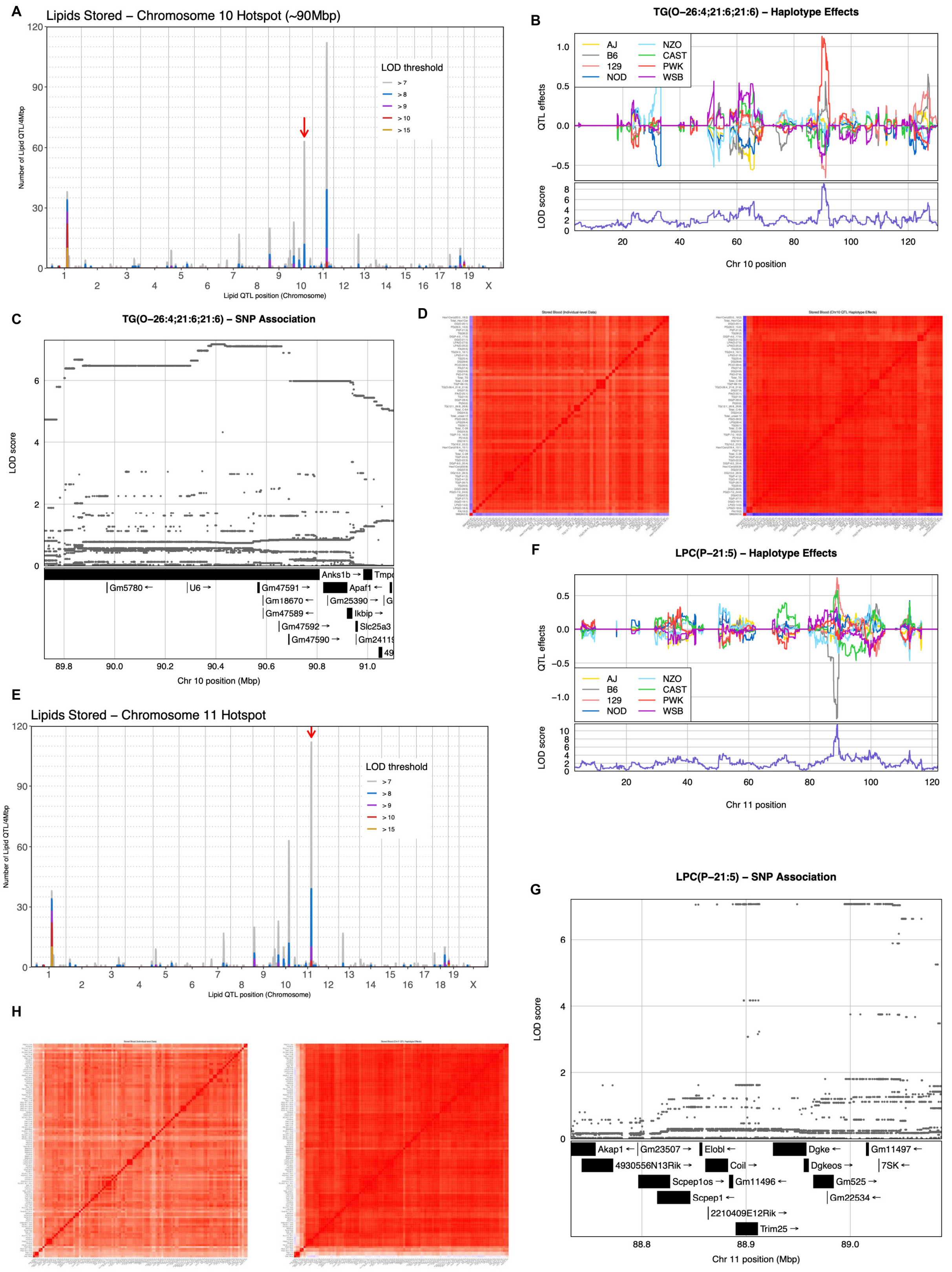


**Supplementary Figure 11 Chromosomes 10 and 11 QTL hotspot are associated with lipid metabolism in mice.** QTL density plot for lipids in stored samples (**A, E**). Genome scan and haplotype effects at the hotspot locus for triacylglycerol

O-26:4;21:6;21:6 [TG(O-26:4;21:6;21:6)] (**B**). SNP associations at TG(O-26:4;21:6;21:6) QTL and genes in the region (**C**). The correlation structure among lipids reflects the QTL hotspot in stored samples (**D**), at both the individual-level data (left) and the QTL haplotype effects (right). Genome scan and haplotype effects at the hotspot locus for lysophosphatidylcholine P-21:5 [LPC(P-21:5)] (**F**). SNP associations at LPC(P-21:5) QTL and genes in the region (**G**). The correlation structure among lipids reflects the QTL hotspot in stored samples (**H**), at both the individual-level data (left) and the QTL haplotype effects (right).

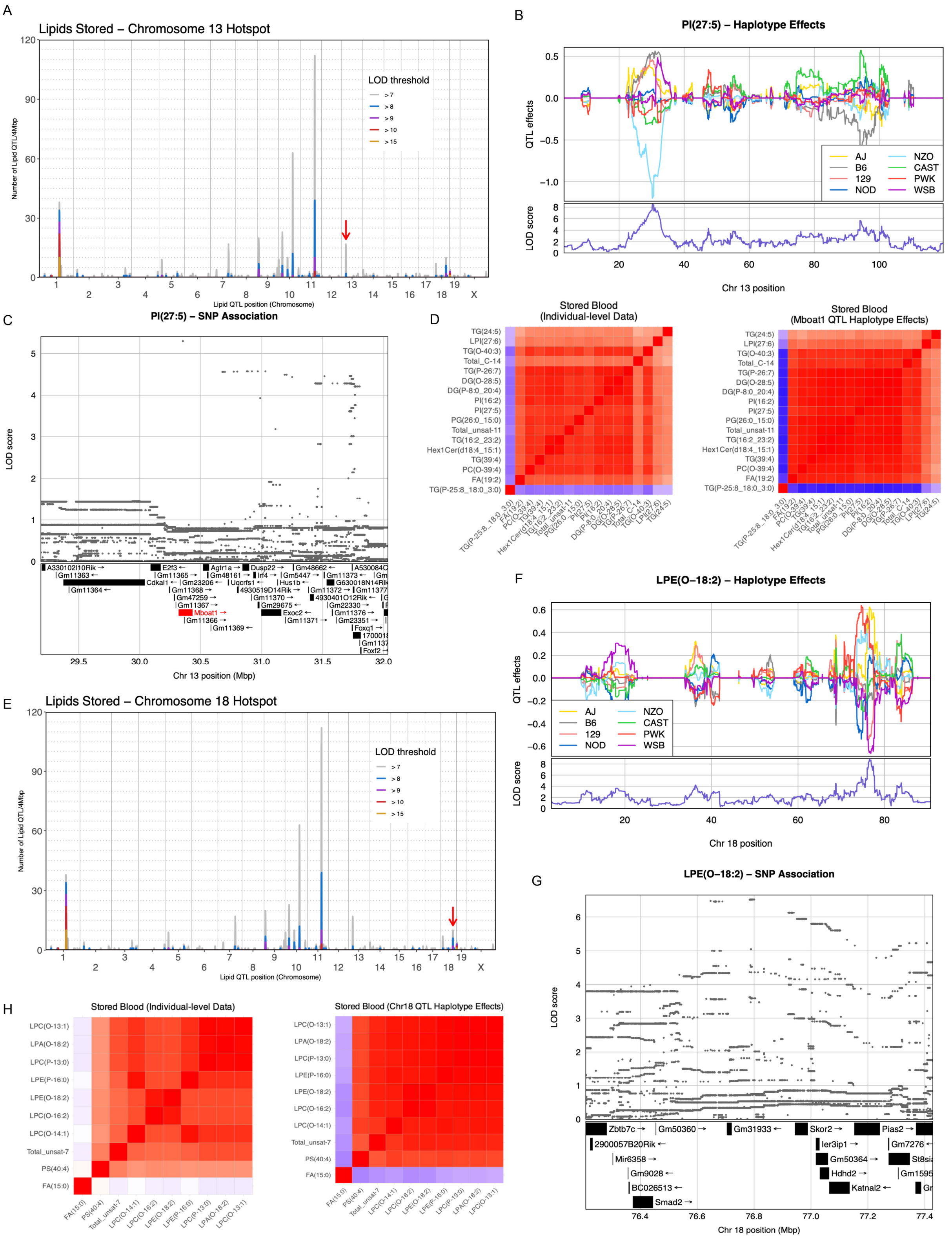


**Supplementary Figure 12 – Chromosome 13 QTL hotspot at the Mboat1 locus and Chromosome 18 QTL hotspot are associated with lipid metabolism in mice.** QTL density plot for lipids in stored samples (**A, E**). Genome scan and haplotype effects at the Mboat1 locus for phosphatidylinositol 27:5 [PI(27:5)] (**B**). SNP associations at PI(27:5) QTL and genes in the region (**C**). The correlation structure among lipids reflects the QTL hotspot in stored samples (**D**), at both the individual-level data (left) and the QTL haplotype effects (right). Genome scan and haplotype effects at the hotspot locus for lysophosphatidylethanolamine O-18:2 [LPE(O-18:2)] (**F**). SNP associations at LPE(O-18:2) QTL and genes in the region (**G**). The correlation structure among lipids reflects the QTL hotspot in stored samples (**H**), at both the individual-level data (left) and the QTL haplotype effects (right).

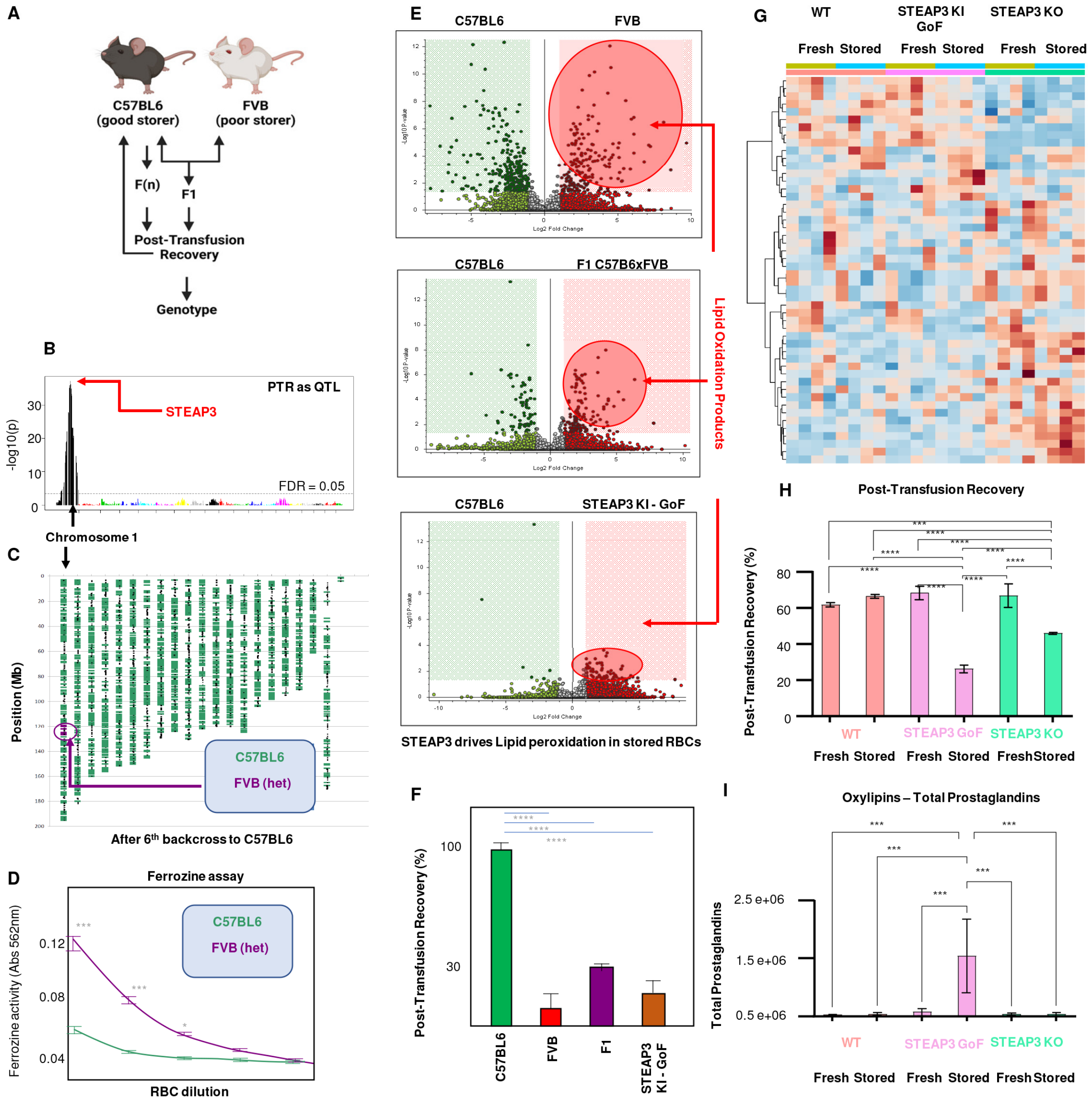


**Supplementary Figure 13 – STEAP3 regulates lipid peroxidation in stored murine RBCs.** Multi-generational crosses of good storer C57BL6/J mice with poor storer FVB mice informed a QTL analysis that identified a region on chromosome 1 coding for the ferrireductase STEAP3 that impacts lipid peroxidation in stored murine RBCs (**A-D**). Knocking in that hypermorphic STEAP3 from FVB mice into C57BL6/J mice increases lipid peroxidation (Volcano plots highlight oxylipins in red - **E**) and decreases post-transfusion recovery in good storing mice (**F**). STEAP3 KO also has a significant impact on RBC metabolism upon storage (**G**), both KO and hypermorphic gain of function (GOF) knock in (KI) resulting in decreases in PTR in mice (**H**). However, only GOF KI mice but not KO show elevated levels of oxylpins in end of storage RBCs (**I**).

**A** List of Common STEAP3 SNPs and prevalence in heterozygosity (1) or homozygosity (2) in the REDS RBC Omics Index Donor population

| SNP | 0 | 1 | 2 |
|---|---|---|---|
| chr2:119983029:T/C:1:rs838108:intronic | 4822 | 5652 | 2042 |
| chr2:119987726:G/C:1:rs708673:intronic | 4914 | 5754 | 1864 |
| chr2:120021917:C/A:1:AX-40642155:3downstream,3utr | 5588 | 5264 | 1609 |
| chr2:120000485:G/T:1:rs60709211:5upstream,intronic | 10551 | 1832 | 90 |
| chr2:120023188:G/A:1:rs4849771:3downstream,3utr | 10858 | 1622 | 69 |
| chr2:120006664:G/C:1:rs2592075:intronic | 10995 | 1513 | 53 |
| chr2:120012412:C/T:1:rs41279770:coding syn | 11139 | 1314 | 52 |
| chr2:120005312:G/A:1:rs17013371:coding nonsyn | 11611 | 887 | 49 |
| chr2:120004946:G/A:1:rs79425886:intronic | 11064 | 1382 | 45 |
| chr2:119982367:A/C:1:rs73948615:intronic | 11998 | 527 | 41 |
| chr2:120005569:C/T:1:rs17805141:coding syn | 11497 | 1021 | 31 |
| chr2:120004079:G/A:1:rs115704924:intronic | 11565 | 959 | 26 |
| chr2:120022977:G/A:1:rs3731602:3downstream,3utr | 12154 | 356 | 18 |
| chr2:119989209:C/T:1:rs80050966:intronic | 12280 | 241 | 11 |
| chr2:119994500:T/C:1:rs113848870:intronic | 12133 | 206 | 11 |
| chr2:120002320:C/T:1:rs111823088:intronic | 12018 | 500 | 9 |
| chr2:119994299:C/T:1:rs111945827:intronic | 12378 | 172 | 8 |
| chr2:120011691:T/C:1:rs115746244:intronic | 12357 | 154 | 5 |
| chr2:120018598:T/C:1:rs115030775:intronic | 12461 | 90 | 3 |
| chr2:120018923:G/A:1:rs77428881:intronic | 12258 | 259 | 3 |
| chr2:120020755:G/A:1:rs61746941:coding syn,3utr | 12480 | 78 | 1 |
| chr2:120005613:G/A:1:rs147820529:coding nonsyn | 12544 | 20 | 1 |
| chr2:119984439:G/T:1:rs115803326:intronic | 12491 | 70 | 1 |
| chr2:120020901:C/T:1:rs141086904:3utr,coding nonsyn | 12508 | 56 | 0 |
| chr2:120003239:T/G:1:rs185132453:coding nonsyn | 12550 | 1 | 0 |
| chr2:120003463:G/A:1:rs181208641:coding nonsyn | 12408 | 16 | 0 |
| chr2:120005676:T/C:1:rs144575753:coding nonsyn | 12553 | 5 | 0 |
| chr2:120005751:G/A:1:rs199836424:coding nonsyn | 12488 | 37 | 0 |
| chr2:120012405:G/A:1:rs41279768:coding nonsyn | 12496 | 53 | 0 |
| chr2:119987140:C/T:1:rs75589170:intronic | 12514 | 49 | 0 |
| chr2:119987437:C/T:1:rs186962782:intronic | 12082 | 465 | 0 |
| chr2:119992184:C/T:1:rs111397350:intronic | 12528 | 37 | 0 |
| chr2:119994065:G/A:1:rs114783565:intronic | 12507 | 52 | 0 |
| chr2:120004155:C/T:1:rs113452357:intronic | 12501 | 43 | 0 |
| chr2:120007568:G/T:1:rs117851232:intronic | 12490 | 56 | 0 |
| chr2:120014236:G/T:1:rs3731604:intronic | 12479 | 63 | 0 |
| chr2:120014716:C/T:1:rs115277200:intronic | 12470 | 46 | 0 |

**B**

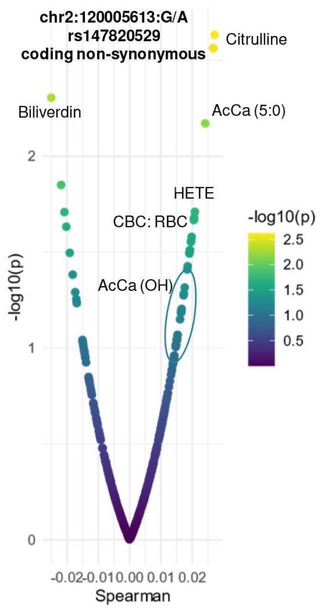


**Supplementary Figure 14 – Allele frequency STEAP3 polymorphisms in the REDS RBC Omics index donor population.** In **A**, 0, 1 and 2 indicate homozygous dominant, heterozygous and homozygous recessive for each STEAP3 SNP in the first column, respectively. In **B**, metabolic correlates to the STEAP3 rs147820529 SNP further confirm the association of non-synonymous coding STEAP3 SNPs to lipid peroxidation products (e.g., HETE).

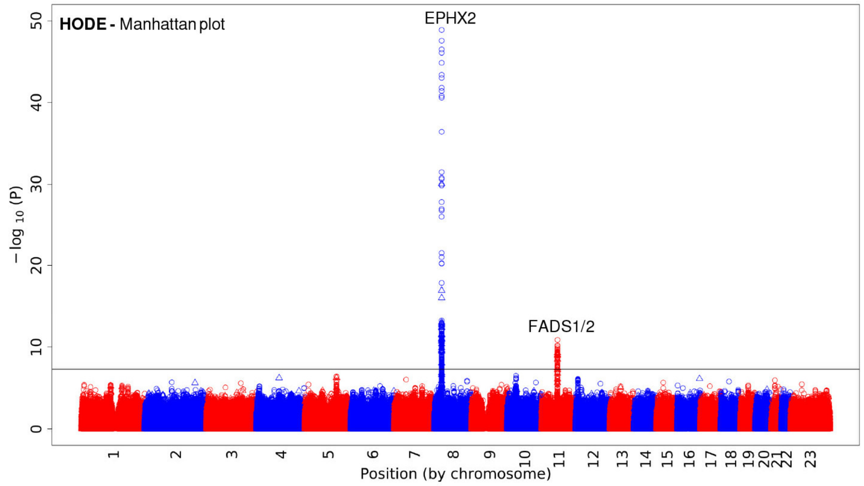


**Supplementary Figure 15 – Manhattan plot for hydroxyoctadecadienoic acid (HODEs – all isomers) in REDS RBC omics.**

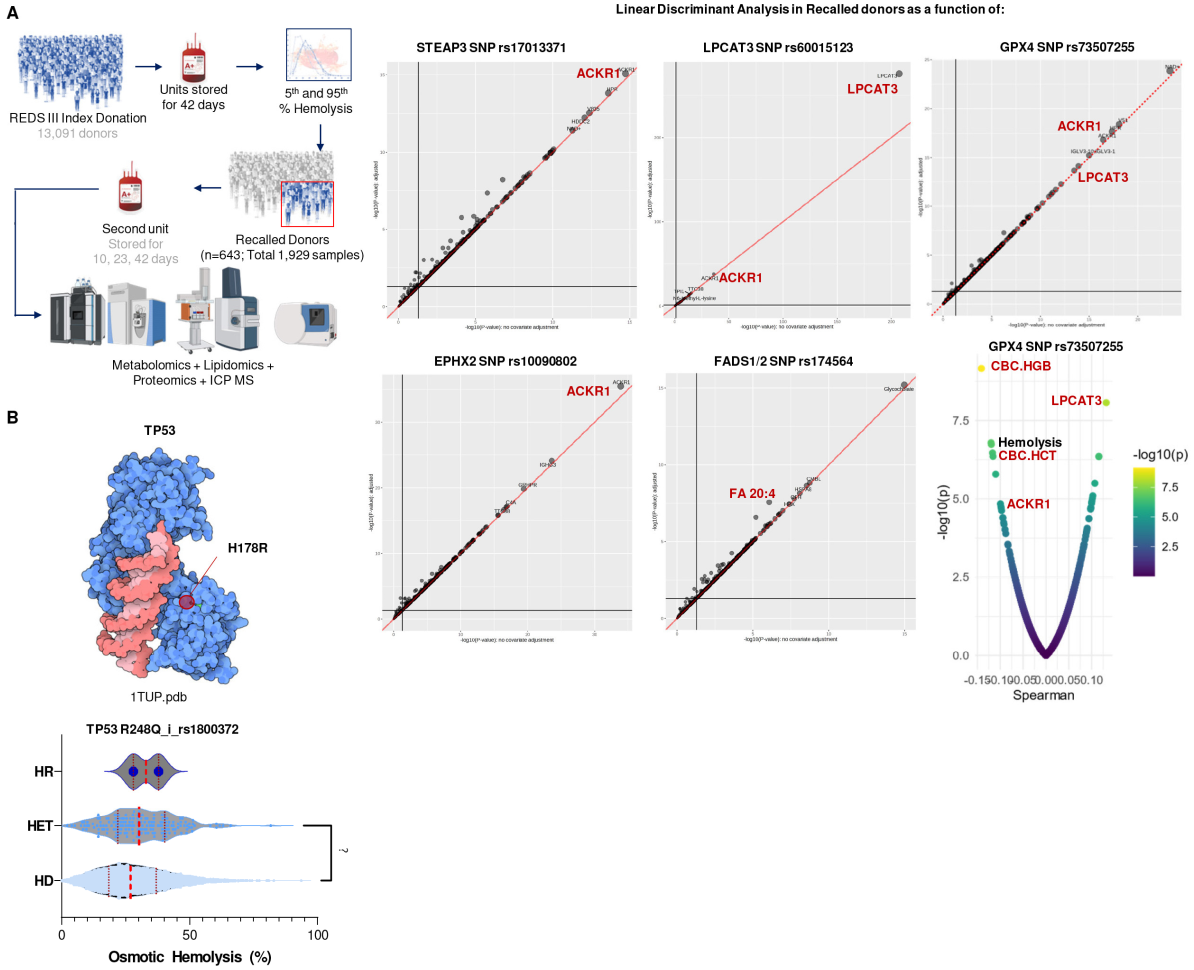


**Supplementary Figure 16 – SNPs in gene associated with oxylipins in stored human RBCs are all also associated to the Duffy atypical chemokine receptor 1 (ACKR1 - A).** In **B**, the TP53 SNP rs1800372 was observed in homozygosity in two blood donors, while more frequently observed in heterozygosity. This potentially pathogenic TP53 SNP is not only associated with the functional R248Q mutation, but here also observed to associate with elevated susceptibility to hemolysis following osmotic insults (? indicates $p<0.05$).
